# Yeast HMGB protein Hmo1 is a multifaceted regulator of DNA damage tolerance

**DOI:** 10.1101/2025.02.21.639549

**Authors:** Jinlong Huo, Anhui Wei, Na Guo, Ruotong Wang, Xin Bi

## Abstract

The chromosome architecture protein Hmo1 in *Saccharomyces cerevisiae* is categorized as an HMGB protein as it consists of two structure-specific DNA binding HMGB motifs. However, it deviates from canonical HMGB proteins that have acidic C-terminal domains (CTDs) by bearing a basic one. Hmo1 plays diverse functions in gene regulation and genome integrity. There is evidence implicating Hmo1 in DNA damage tolerance (DDT) that enables DNA replication to bypass lesions on the template. Hmo1 is believed to direct DNA lesions to the error-free template switching (TS) pathway of DDT and aids in the formation of the key TS intermediate sister chromatid junction (SCJ), but the underlying mechanism(s) has yet to be resolved. In this work, we took genetics and molecular biology approaches to further investigate the role of Hmo1 in DDT. We found extensive functional interactions of Hmo1 with components of the genome integrity network in cellular response to the genotoxin methyl methanesulfonate (MMS), implicating Hmo1 in multiple processes in the execution or regulation of homology-directed DNA repair, replication-coupled chromatin assembly, and cell cycle checkpoint. Notably, our data pointed to a role of Hmo1 in directing SJC to the nuclease-mediated *resolution* pathway instead of the helicase/topoisomerase mediated *dissolution* pathway for processing/removal. They also suggested that Hmo1 modulates both the recycling of parental histones and the deposition of newly synthesized histones at the replication fork to ensure proper chromatin formation on nascent DNA that is important for DDT. We found evidence that Hmo1 antagonizes the function of histone H2A variant H2A.Z (also known as Htz1 in yeast) in DDT. We also found that Hmo1 is required for DNA damage checkpoint signaling induced by MMS as wells as for DNA negative supercoiling as a proxy of chromatin structure. Moreover, we obtained evidence indicating that the CTD of Hmo1 may or may not be required for its function in DDT depending on the genetic background of the host. Taken together, our findings demonstrate that Hmo1 can contribute to, or regulate, multiple processes of DDT via different mechanisms.

## Introduction

Chromatin plays a critical role in the maintenance and function of eukaryotic genome by regulating DNA transactions including replication, recombination, and repair as well as gene transcription. The basic unit of chromatin is the nucleosome composed of 147 base pairs of DNA wrapping around a protein core made of two copies of each of histones H2A, H2B, H3, and H4 (Luger et al. 1997). Nucleosomes are connected by linker DNA sequences to form the primary chromatin structure. Formation of higher order chromatin structure is facilitated by linker histone H1 that associates with linker DNA (Hergeth and Schneider 2015). Chromatin compacts and protects DNA but also hinders DNA transactions. Cells have evolved mechanisms to modify/remodel chromatin to create global or local chromatin states/environments that are suitable for genome functions. One such mechanism is the modulation of chromatin dynamics by the high mobility group (HMG) proteins, an evolutionally conserved group of non-histone chromatin architectural proteins (Stros 2010). HMG proteins comprise the HMBA, HMGB and HMGN families, of which HMGB family is by far the largest. An HMGB protein typically consists of one or more HMG-boxes (aka HMGB motifs) and an acidic carboxyterminal domain (CTD) containing consecutive glutamate and aspartate residues. A HMGB motif is a sequence-nonspecific DNA binding module that recognizes altered DNA structures such as hemicatenane and four-way junction and can distort DNA via bending and unwinding (Stros 2010). The CTD of an HMGB protein regulates its DNA binding properties (Stros 2010). In mammals, HMGB proteins are known to interact with nucleosomes to evict linker histone H1, making chromatin more accessible to other chromatin binding factors (Postnikov and Bustin 2016).

*Saccharomyces cerevisiae* Hmo1 is deemed an HMGB protein as it contains two HMGB motifs that binds DNA (Lu et al. 1996; Panday and Grove 2017). However, unlike canonical HMGB proteins, Hmo1 has a basic, lysine rich, CTD, similarly as linker histone H1. In line with its special domain structure, Hmo1 exhibits functional characteristics of both HMGB proteins and linker histone H1 (Panday and Grove 2017; Bi 2024). On one hand, like HMGB proteins, Hmo1 destabilize nucleosomes (McCauley et al. 2019; Malinina et al. 2022) and facilitates the recruitment of gene regulators (Hepp et al. 2017; Amigo et al. 2022). On the other hand, like linker histones, Hmo1 makes yeast genome refractory to nuclease digestion possibly by promoting chromatin compaction and rendering it less dynamic (Lu et al. 1996; Panday and Grove 2016). Notably, this function of Hmo1 is dependent on its CTD (Panday and Grove 2016). What determines whether Hmo1 performs HMGB-like or linker histone-like functions in the cell is not clear.

As an abundant protein that associates with chromatin throughout the cell cycle (Bermejo et al. 2009), Hmo1 impacts multiple cellular functions (reviewed in Bi 2024). Hmo1 is important for cell growth as cells lacking Hmo1 exhibit a severe slow growth phenotype due to a significant lengthening of the doubling time/cell cycle (Lu et al.1996). Hmo1 acts as a transcription factor to promote the expression of genes involved in ribosome biogenesis (Bi 2024). Hmo1 was recently found to bind the boundaries of RNA polymerase II transcribed genes and help establish a shared topological configuration of these genes (Achar et al. 2020). Hmo1 was also found to affect the repair of double stranded DNA (dsDNA) breaks (DSBs) by stabilizing chromatin and hindering DNA end resection (Panday et al. 2015; Panday et al. 2017). In addition, there is evidence that Hmo1 regulates cellular tolerance *to* methyl methanesulfonate (MMS), a genotoxin that methylates DNA and stalls DNA replication (Gonzalez-Huici et al. 2014). During DNA replication, damages on the template may render leading strand and lagging strand synthesis discontinuous due to the uncoupling of DNA polymerase from DNA helicase or re-initiation of DNA synthesis at a distance from the lesion (Sogo et al. 2002; Byun et al. 2005). This results in single stranded DNA (ssDNA) gaps in the wake of stalled replication forks which can be repaired/filled by two mechanistically distinct DDT pathways that enable the replisome to bypass DNA lesions and complete DNA replication (reviewed in Bi 2015 and Branzei and Psakhye 2016). One mechanism is translesion synthesis (TLS) in which the replicative DNA polymerase is temporarily replaced by a specialized TLS polymerase capable of replicating across DNA lesions (Fig. 1). The other is template switching (TS) which involves the switching of the stalled nascent DNA strand from damaged template to undamaged sister strand for extension past the lesion (Fig. 1). Initiation of TS is accomplished by the invasion of the sister chromatid by the stalled strand (Fig. 1). The resulting D-loop is subsequently converted into a joint molecule called sister chromatid junction (SCJ) after DNA synthesis using the intact sister strand as template (Fig. 1). SCJ can be disentangled to yield two duplex DNA strands via dissolution by the helicase/topoisomerase complex Sgs1/Top3/Rmi1 (STR) or resolution by the structure-specific nuclease Mus81/Mms4 complex (Fig. 1). Note although TLS is mechanistically simple, it is error-prone and largely accountable for mutagenesis. TS, on the other hand, is complex but error-free, and is therefore believed to be the preferred pathway. How cells choose a DTT pathway in response to genotoxin-induced DNA lesions is not clear. Gonzalez-Huici et al. found evidence suggesting that Hmo1 helps to channel MMS-induced DNA lesions to the TS pathway and facilitates the formation of SCJ (Gonzalez-Huici et al. 2014), but the underlying mechanism has yet to be resolved.

**Fig. 1.**
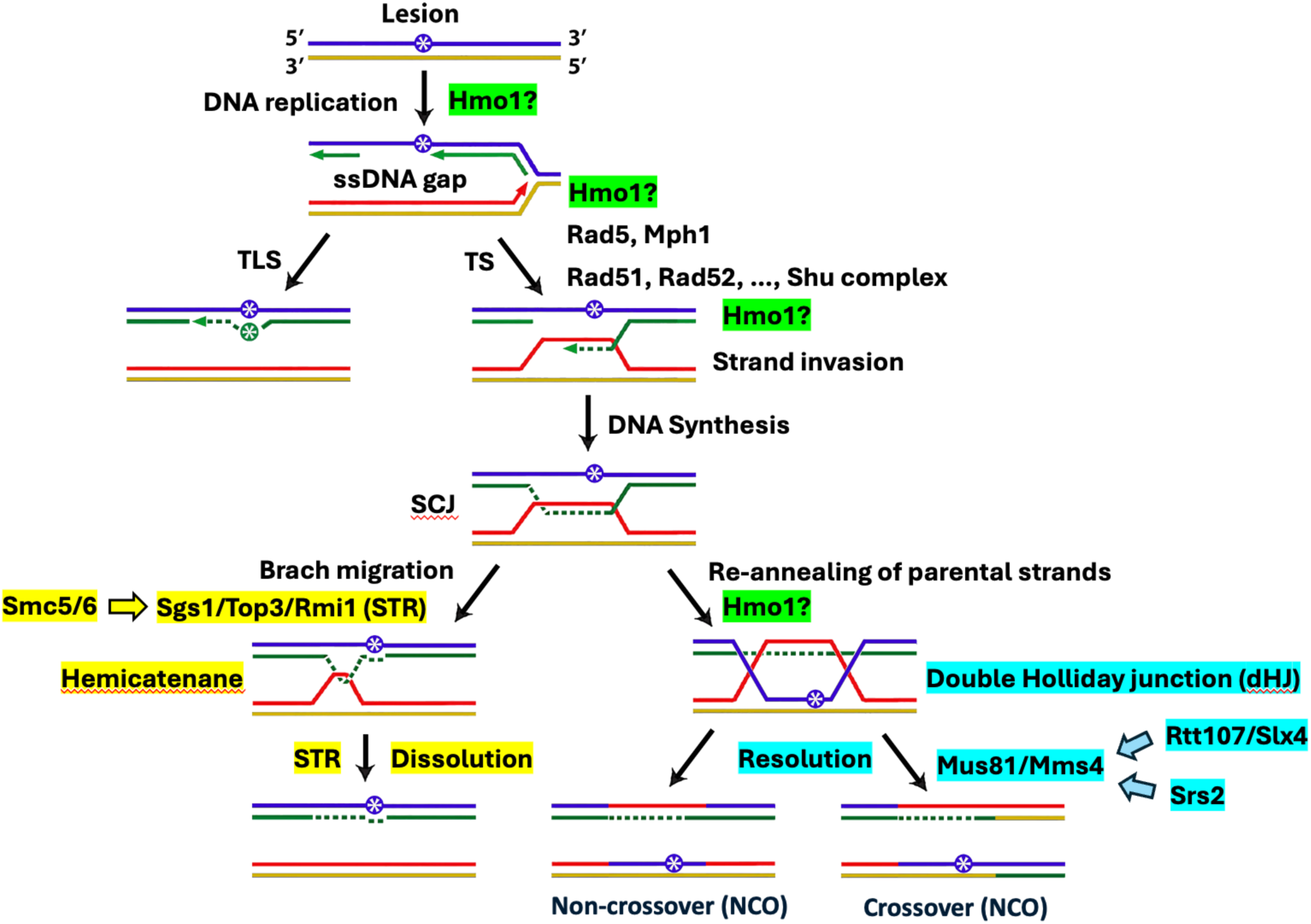
Mechanisms of DNA damage tolerance (DDT). DNA lesion (asterisk in blue circle) inflicted by genotoxins may stall DNA replication. Uncoupling of DNA polymerase and helicase or reinitiation of DNA synthesis at a distance leads to the formation of ssDNA gaps on leading and lagging strands (a ssDNA gap on lagging strand is shown). Repair/filling of ssDNA gaps can be achieved via the translesion systhesis (TLS) or template switching (TS) pathways that are mechanistically different. SCJ, sister chromatid junction; “Hmo1?” denotes possible position/step of the DDT mechanisms where Hmo1 may act or regulate. See the text for descriptions.

In this work, we took genetics and molecular biology approaches to further investigate the role of Hmo1 in DDT. We found extensive functional interactions of Hmo1 with components of the genome integrity network in cellular response to MMS, implicating Hmo1 in multiple processes in the execution or regulation of homology-directed DNA repair, replication-coupled chromatin assembly, and cell cycle checkpoint that participate in, or impact, DDT. Notably, our data pointed to a role of Hmo1 in directing TS intermediate SJC to Mus81/Mms4-mediated *resolution* pathway instead of STR-mediated *dissolution* pathway for processing/removal. They also suggested that Hmo1 modulates both the recycling of parental histones and the deposition of newly synthesized histones at the replication fork that are important for DDT. We found evidence that Hmo1 antagonizes histone H2A variant H2A.Z (also known as Htz1 in yeast) in DDT. We also found that Hmo1 is required for DNA damage checkpoint signaling induced by MMS as wells as DNA negative supercoiling as a proxy of chromatin structure. Moreover, we showed that the CTD of Hmo1, and by inference its DNA bending activity, is dispensable for its function in promoting checkpoint or DNA supercoiling. Interestingly, we obtained evidence indicating that the CTD of Hmo1 may or may not be required for its function in DDT depending on the genetic background of the host. Taken together, our findings suggest that Hmo1 can contribute to, or regulate, DDT via multiple mechanisms.

## Materials and Methods

### Yeast strains

Yeast strains used in this work are listed in Table 1. Gene deletion was done by replacing the ORF of the gene of interest with *NatMX* or *URA3*, which was verified by Southern blotting or PCR. Strains W1588-4C and T382-P4 were obtained from Dr. Xiaolan Zhao (Memorial Sloan Kettering Cancer Center), QY364 and QY375 from Dr. Stephen Kron (University of Chicago), PY39, PY39-8, and PY30-79 from Dr. Peter Burgers (Washington University). Plasmids pRS416-*HMO1* (pHWT) and pRS416-*hmo1-AB* (pHAB) were obtained from Dr. Anne Grove (Louisiana State University). Synthetic complete (SC) medium was used for growing yeast cells.

**Table 1.**
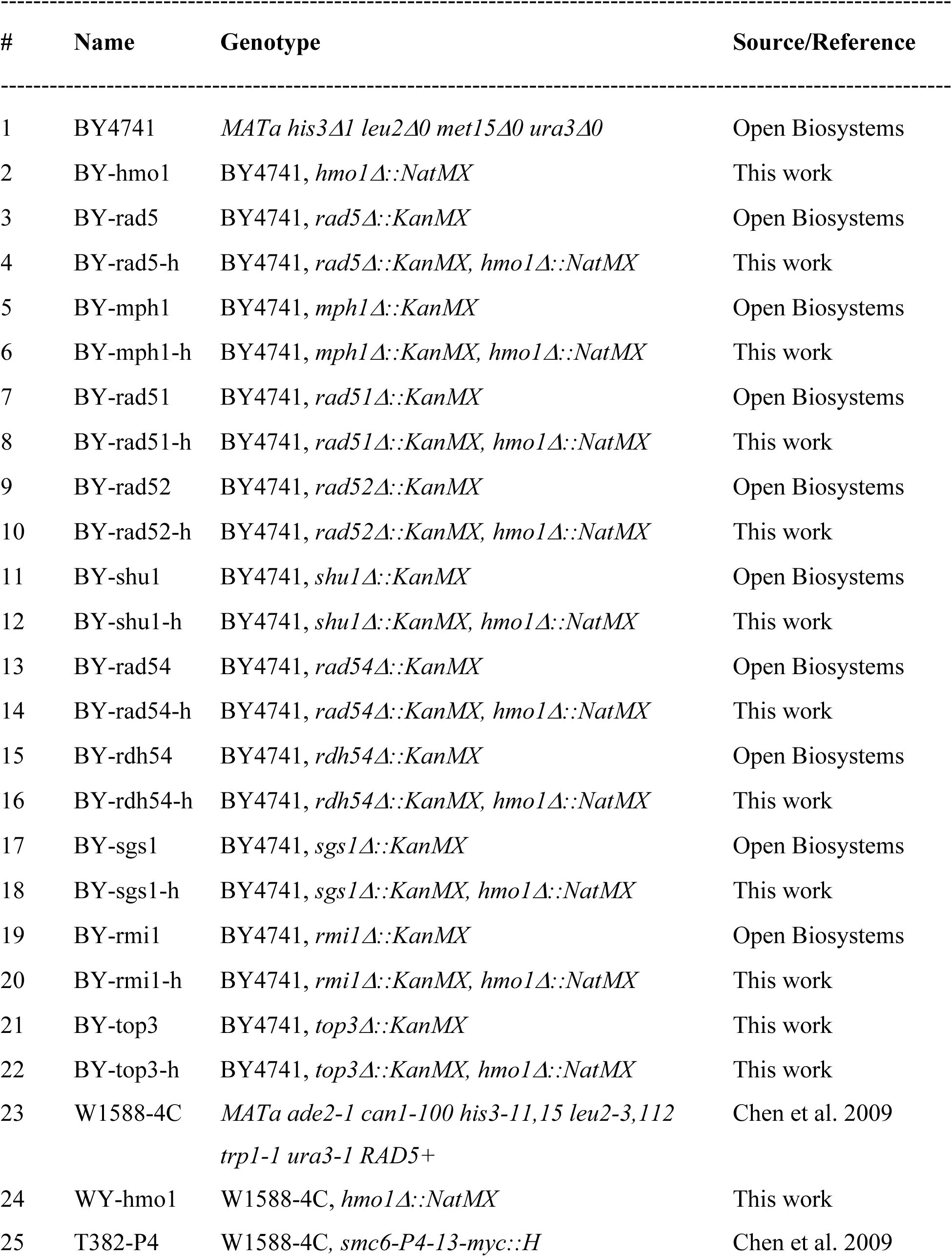
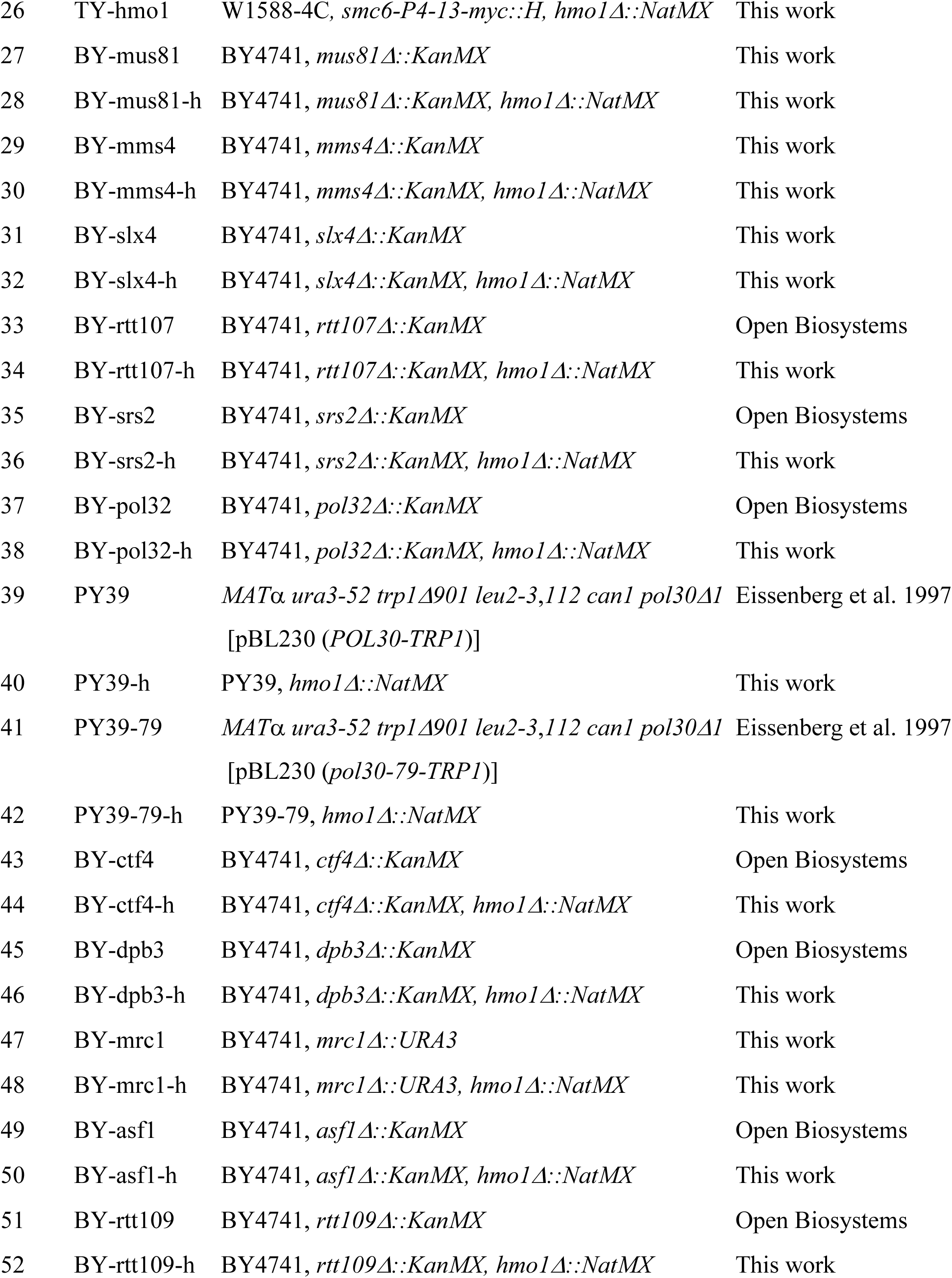
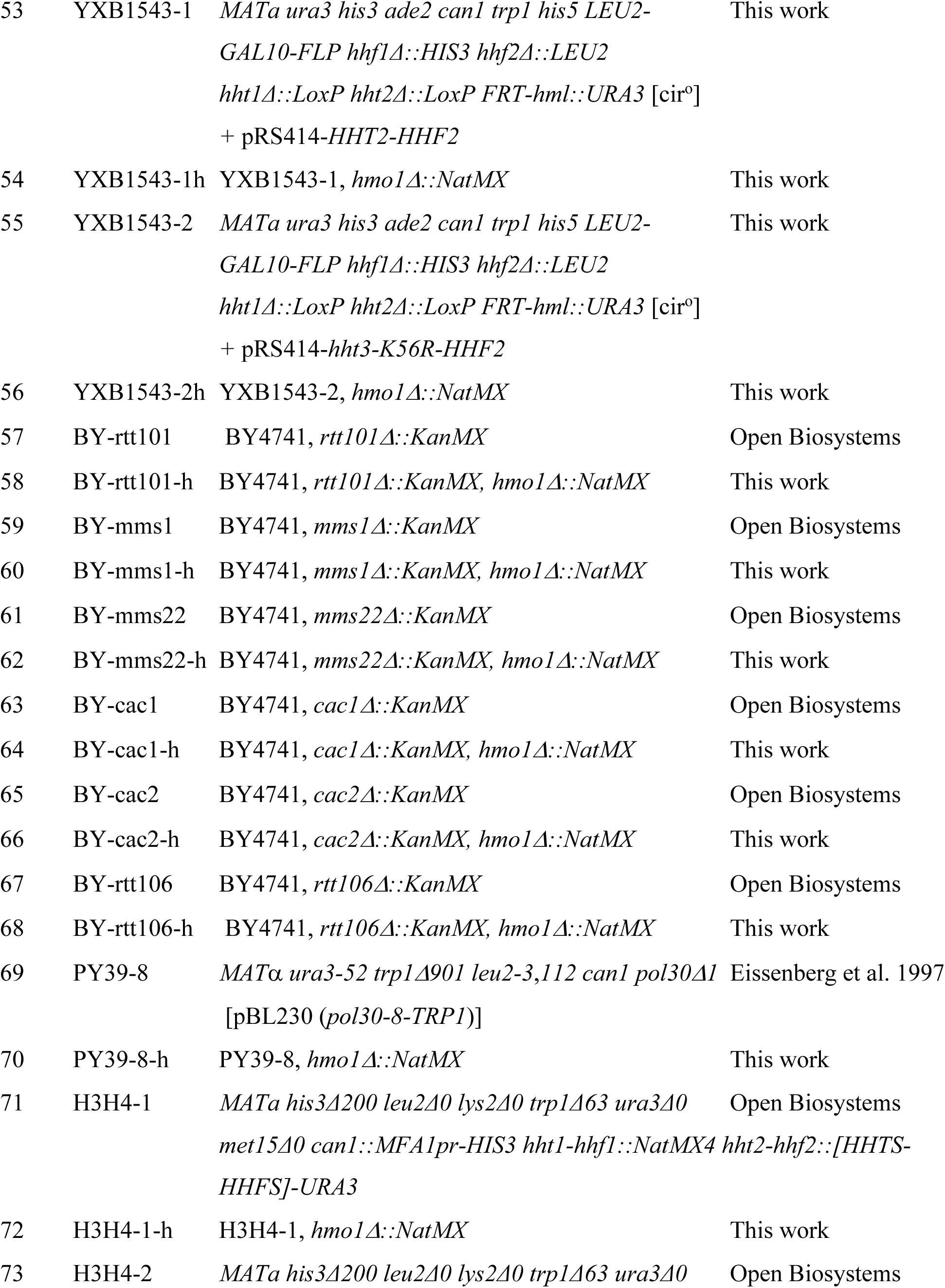
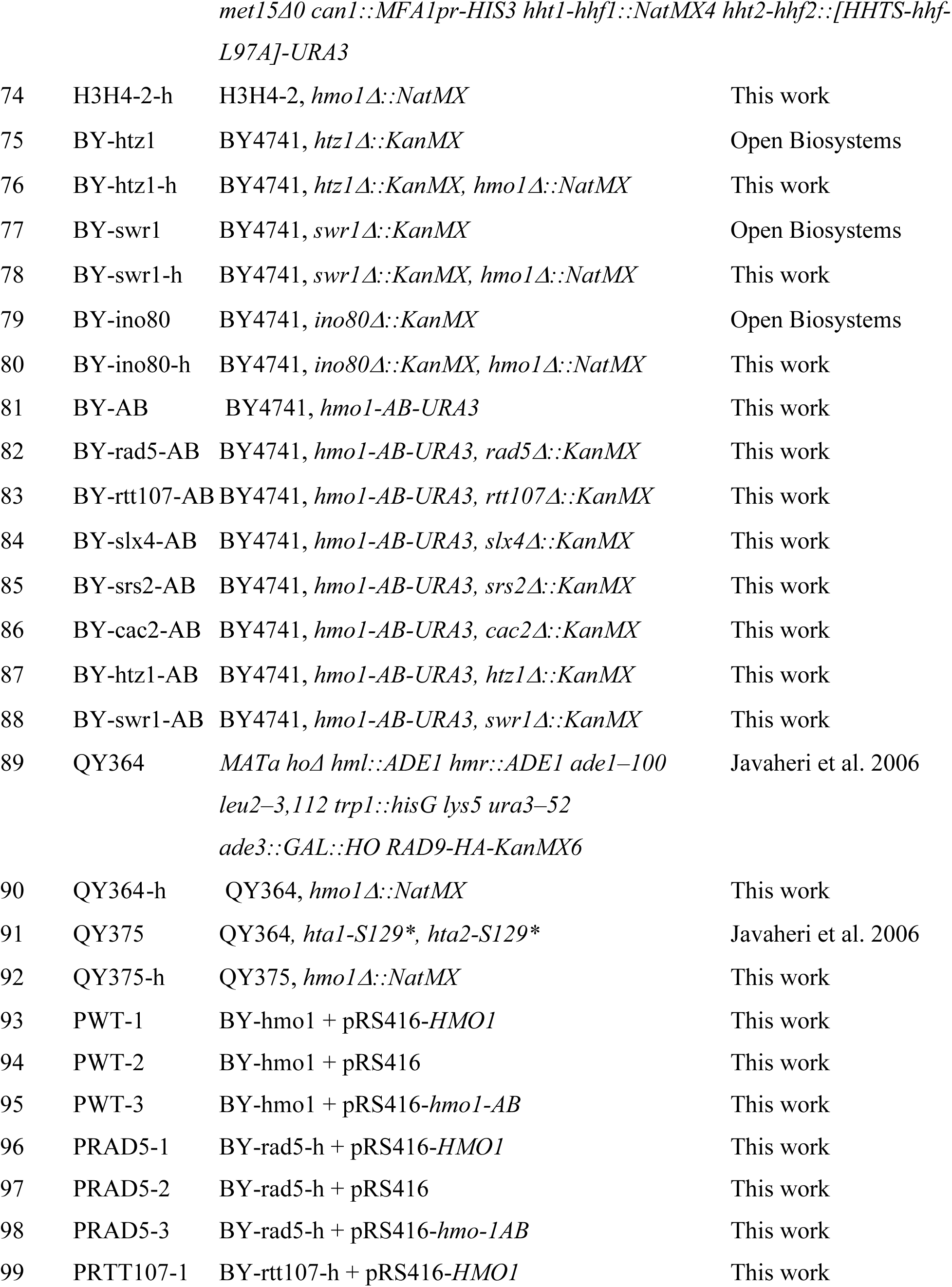
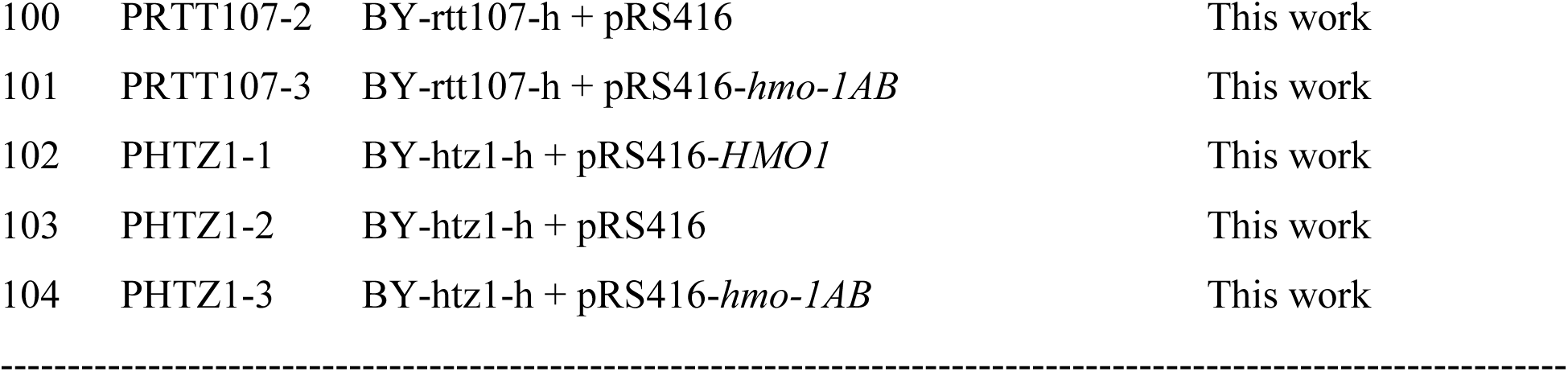
Yeast strains.

### SDS-PAGE and Western blotting

Protein extracts from yeast cells were obtained by TCA extraction. Ten μg of proteins from each sample of cells was electrophoresed by SDS-PAGE in 4-12% gradient gels. Western blotting was done using LI-COR Odyssey CLx Infrared Imaging System (LI-COR Biosciences). Antibodies used were: goat polyclonal anti-Rad53 (yC-19: sc6749, Santa Cruz Biotechnology) and rabbit polyclonal anti-HA (H6908, Sigma-Aldrich). Secondary antibodies used were LI-COR IRDye 800CW goat polyclonal anti-rabbit IgG (H+L) 926-32211 and LI-COR IRDye 800CW donkey anti-goat IgG (H+L) 926-32214.

### Analysis of the topology of yeast 2-micron (2μ) plasmid

Yeast cells were grown in liquid SC medium to late log phase. Nucleic acids were isolated from yeast cells using the glass bead method and fractionated on 0.8% agarose gel in 0.5 x TPE (45 mM Tris, 45 mM phosphate, 1 mM EDTA, pH 8.0) supplemented with 12 μg/ml chloroquine. After Southern blotting, topoisomers of 2μ plasmid were detected by hybridization with a radioactively labeled fragment of 2μ plasmid as the probe. Under the electrophoresis conditions used in this work, more negatively supercoiled topoisomers migrate more slowly.

## Results

### Hmo1 directs SCJ intermediate of DDT to Mus81/Mms4 nuclease-mediated resolution pathway

DDT, also known as post-replication repair, is intimately intertwined with DNA replication (Branzei and Psakhye 2016). The sliding DNA clamp PCNA, a key replication factor, plays a central role in controlling the choice of DDT pathways (Moldovan et al. 2007; Branzei and Psakhye 2016). PCNA mono-ubiquitination promotes TLS whereas PCNA polyubiquitination by Rad5/Ubc13/Mms2 activates the TS pathway of DDT. Rad5 also has DNA helicase activity that promotes fork reversal that can lead to SCJ formation at a stalled replication fork (Blastyak et al. 2007; Meng and Zhao 2017). Hmo1 has been implicated in TS pathway of DDT as deletion of the *HMO1* gene (*hmo1Δ*) suppresses hypersensitivity of *rad5Δ* mutant to MMS (Gonzalez-Huici et al. 2014) (Fig. 2). Like Rad5, the Mph1 helicase also promotes fork reversal at a stalled replication fork (Meng and Zhao 2017). We found that *hmo1Δ* also suppressed MMS-sensitivity of *mph1Δ* mutant (Fig. 2), indicating that Hmo1 functions upstream of Rad5 and Mph1 to possibly channel DNA lesions inflicted by MMS to TS pathway. Gonzalez-Huici et al. found evidence that Hmo1 aids in the formation of SCJ destinated to be processed by TS (Gonzalez-Huici et al. 2014) (Fig. 1). It was believed that the lack of TS in *rad5Δ* mutant led to the toxic accumulation of SCJs, causing MMS-hypersensitivity, and additional deletion of Hmo1would reduce the level of SCJs, thereby mitigating MMS-sensitivity of *rad5Δ* mutant (Gonzalez-Huici et al. 2014). This notion can also explain the suppression of MMS-sensitivity of *mph1Δ* mutant by *hmo1Δ*.

**Fig. 2.**
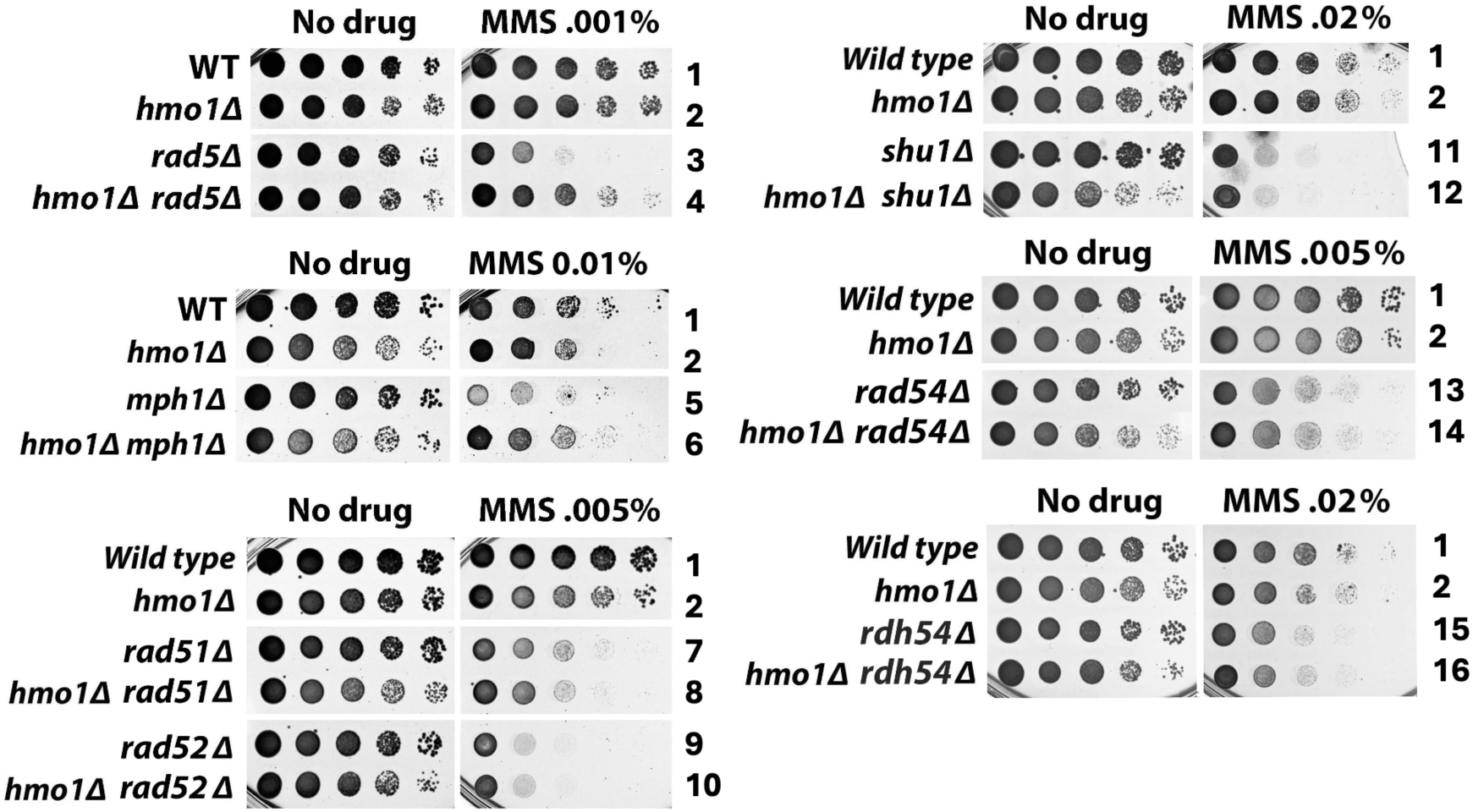
Functional interplay of Hmo1 with Rad5, Mph1 and homologous recombination factors in DDT. Shown are growth phenotypes of indicated strains (#1-16 in **Table 1**) in the presence or absence of MMS. Cells of each strain were grown to saturation, and 10-fold serial dilutions of the culture were spotted on SC (synthetic complete) medium/plate with (No drug) or without MMS. The plates were incubated for 3 days at 30°C before their images were taken.

Given that Hmo1 binds DNA with specificities for altered structures such as hemicatenane and four-way junction that exist in TS intermediates such as SCJ (Fig. 1), we wondered whether Hmo1 is also involved in TS processes downstream of Rad5 and Mph1. Accordingly, we examined the interplay of Hmo1 with other TS factors involved in the formation and processing/removal of SCJ in cellular tolerance to MMS. SCJ formation is mediated by core homologous recombination (HR) factors including the Rad52 epistasis group of proteins and specialized HR factors such as the Shu complex (Shu1/Shu2/Psy3/Csm2) (reviewed in Bonner and Zhao 2016; Martino and Bernstein 2016) (Fig. 1). Rad51 recombinase promotes the invasion of sister chromatid by ssDNA, which is facilitated by Rad52 and Shu complex (Gaines et al. 2015; Godin et al. 2016; Bonner and Zhao 2016; Martino and Bernstein 2016). Rad54 and Rdh54 aid in homology search by Rad51 presynaptic complex (Crickard et al. 2020). We found that *RDA51*, *RAD52*, *SHU1*, *RAD54* and *RDH54* were all epistatic to *HMO1* in MMS-tolerance (Fig. 2), suggesting a role of Hmo1 in TS pathway downstream of SCJ formation mediated by HR factors. To test this notion, we examined if Hmo1 was involved in the processing/removal of SCJ.

SCJ can be converted to a hemicatenane that can be dissolved by the STR helicase/topoisomerase complex, or to a double Holliday junction (dHJ) that can be resolved by the structure-specific nuclease Mus81/Mms4 (reviewed in Bonner and Zhao 2016) (Fig. 1). STR mediated SCJ dissolution yields non-crossover products, whereas Mus81/Mms4 mediated resolution may generate non-crossover and crossover products (reviewed in Bizard and Hickson 2014; Bonner and Zhao 2016) (Fig. 1). We found that *hmo1Δ* exacerbated MMS-sensitivity of *sgs1Δ*, *top3Δ* and *rmi1Δ* mutants (Fig. 3). STR activity is stimulated by Smc5/6-mediated SUMOylation of Sgs1, Top3 and Rmi1 (Bonner et al. 2016). We showed that *hmo1Δ* also exacerbated MMS-sensitivity of cells carrying the *smc6-P4* mutation which disrupts the SUMO ligase function of Smc5/Smc6 (Fig. 3). These results indicated that Hmo1 functions separately from STR in DDT. On the other hand, we showed that *hmo1Δ* suppressed MMS-sensitivity of *mus81Δ* and *mms4Δ* mutants (Fig. 3). Slx4 in complex with Rtt107 is involved in the activation of the nuclease activity of Mus81/Mms4, whereas Srs2 associates with Mus81/Mms4 and promotes its function in SCJ resolution (Gritenaite et al. 2014; Chavdarova et al. 2015). We found *hmo1Δ* to also suppress MMS-sensitivity of *slx4Δ, rtt107Δ* and *srs2Δ* mutants (Fig. 3). These results suggest that Hmo1 acts to direct SCJ to the resolution pathway mediated by Mus81/Mms4, in addition to its function early in DDT to channel DNA lesions into the TS pathway.

**Fig. 3.**
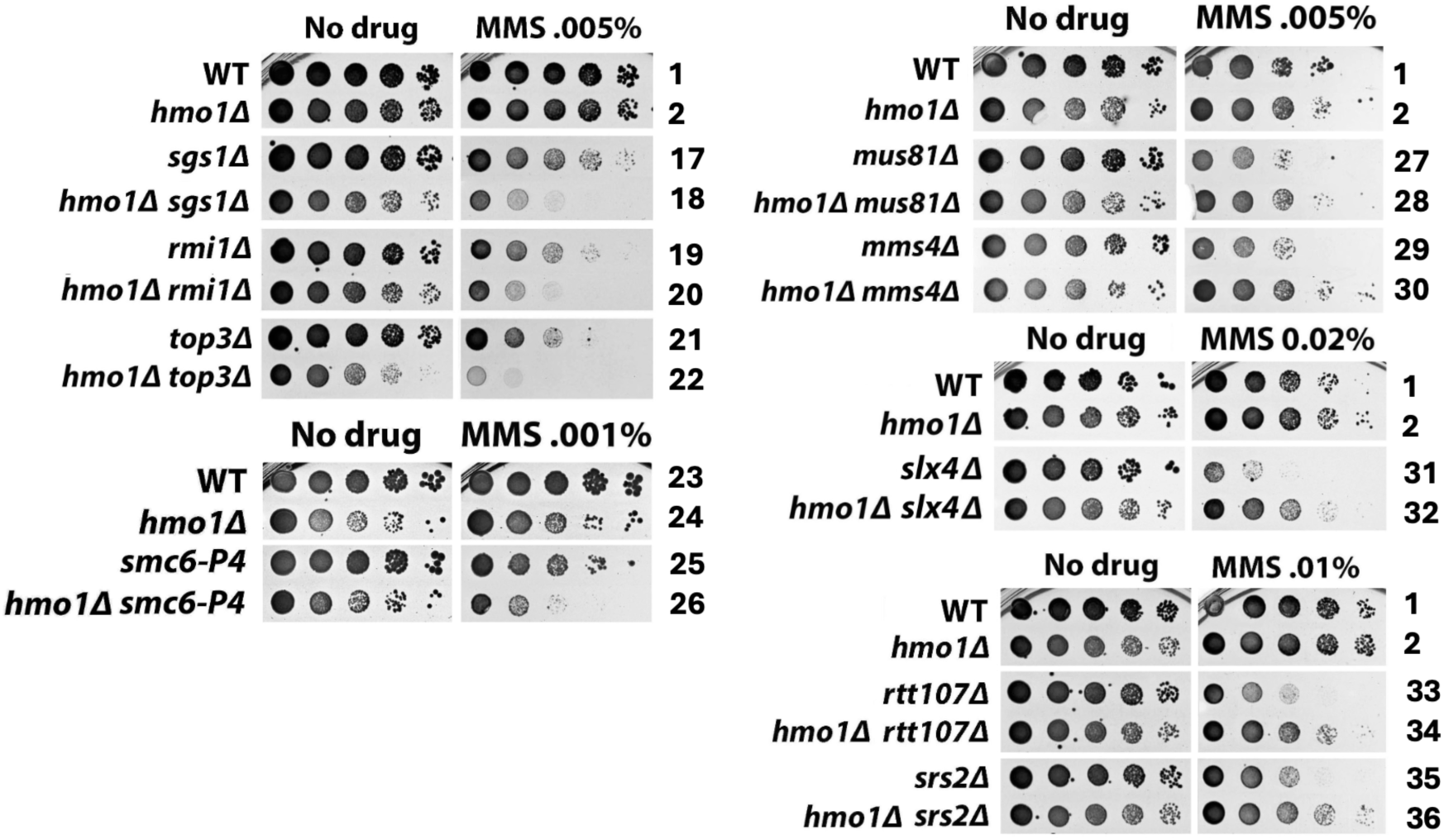
Hmo1 functions in the Mus81/Mms4-mediated resolution pathway independently of the Sgs1/Top3/Rmi1 (STR)-mediated dissolution pathway in DDT. Shown are growth phenotypes of indicated strains in the presence or absence of MMS.

### Hmo1 can contribute to DDT by functioning in DNA replication-coupled chromatin assembly

DNA replication-coupled chromatin assembly plays a critical role in DDT (Hammond-Martel et al. 2021). Given that Hmo1 associates with chromatin throughout the cell cycle (Bermejo et al. 2009) and that Hmo1 can affect nucleosome structure and chromatin compaction (Lu et al. 1996; McCauley et al. 2019; Malinina et al. 2022), it is possible that Hmo1 can contribute to DDT by impacting replication-coupled chromatin assembly. Nucleosome formation on nascent DNA is accomplished by the recycling of parental histones mediated by the replisome and the deposition of newly synthesized histones by chromatin assembly factors (Serra-Cardona and Zhang 2018). At the replication fork, parental histone H3/H4 tetramer is displaced from the nucleosome and then distributed onto both leading and lagging strands by distinct strand-specific mechanisms. Transfer to leading strand is promoted by DNA polymerase Pol ε (Yu et al. 2018; Bellelli et al. 2018) and transfer to lagging strand is governed by the PCNA-Pol δ and Mcm2-Ctf4-Pol α axes (Gan et al. 2018; Tian et al. 2024; Shi et al. 2024) (Fig. 4A). The nonessential Dpb3/Dpb4 subunit of Pol ε and Pol32 subunit of Pol δ both bind histones H3/H4 directly (Tian et al. 2024; Shi et al. 2024). Moreover, the replisome component Mrc1 also plays a role in symmetric distribution of parental histones to sister chromatids (Charlton et al. 2024; Yu et al. 2024; Toda et al. 2024) (Fig. 4A). Newly made histones H3/H4 are deposited to nascent DNA via a histone modification and nucleosome assembly pathway (Collins et al. 2007; Serra-Cardona and Zhang 2017) (Fig. 5A). In this pathway, H3/H4 dimer associates with histone chaperone Asf1 followed by the acetylation of lysine 56 of H3 (H3-K56-Ac) by histone acetylase Rtt109 and subsequent H3 ubiquitination (on K121, K122 and K125) by the Rtt101/Mms1/Mms22 ubiquitinase (Han et al. 2013; Serra-Cardona and Zhang 2017; Hammond-Martel et al. 2021). Histone H3 acetylation and ubiquitination facilitate the transfer of H3/H4 dimer from Asf1 to chromatin assembly factors CAF-1 (Cac1/Cac2/Cca3 complex) and Rtt106 that deposit H3/H4 on nascent strand. Defects in parental histone recycling or new histone deposition make cells sensitive to genotoxins (Hammond-Martel et al. 2021). For instance, the *pol32Δ, ctf4Δ,* or *mrc1Δ* mutation sensitizes cells to MMS (Fig. 4B and 4C). Deletion of any component of new histone deposition pathway or changing H3-K56 to non-acetylable arginine (H3-K56R) also makes cells sensitive to MMS (Fig. 5B and 5C).

**Fig. 4.**
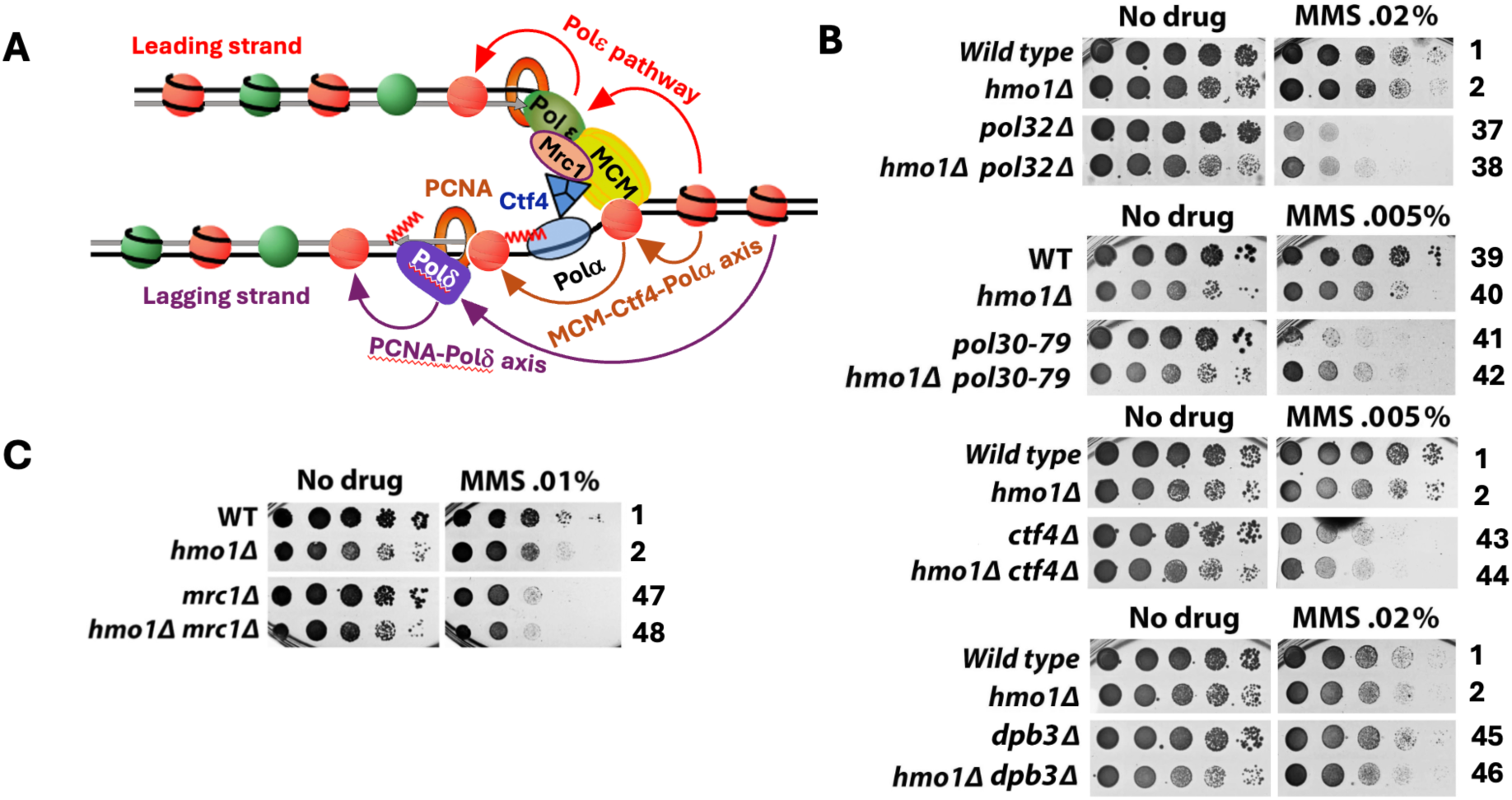
Hmo1 can contribute to DDT by modulating parental histone recycling at the replication fork. (A). Mechanisms of strand-specific transfer of parental histones H3/H4 to leading and lagging strands behind the replication fork. Wiggly line, RNA primer. Green and red circles, nucleosomes bearing new and old H3/H4, respectively. See the text for descriptions. (B) and (C). Growth phenotypes of indicated strains in the presence or absence of MMS.

**Fig. 5.**
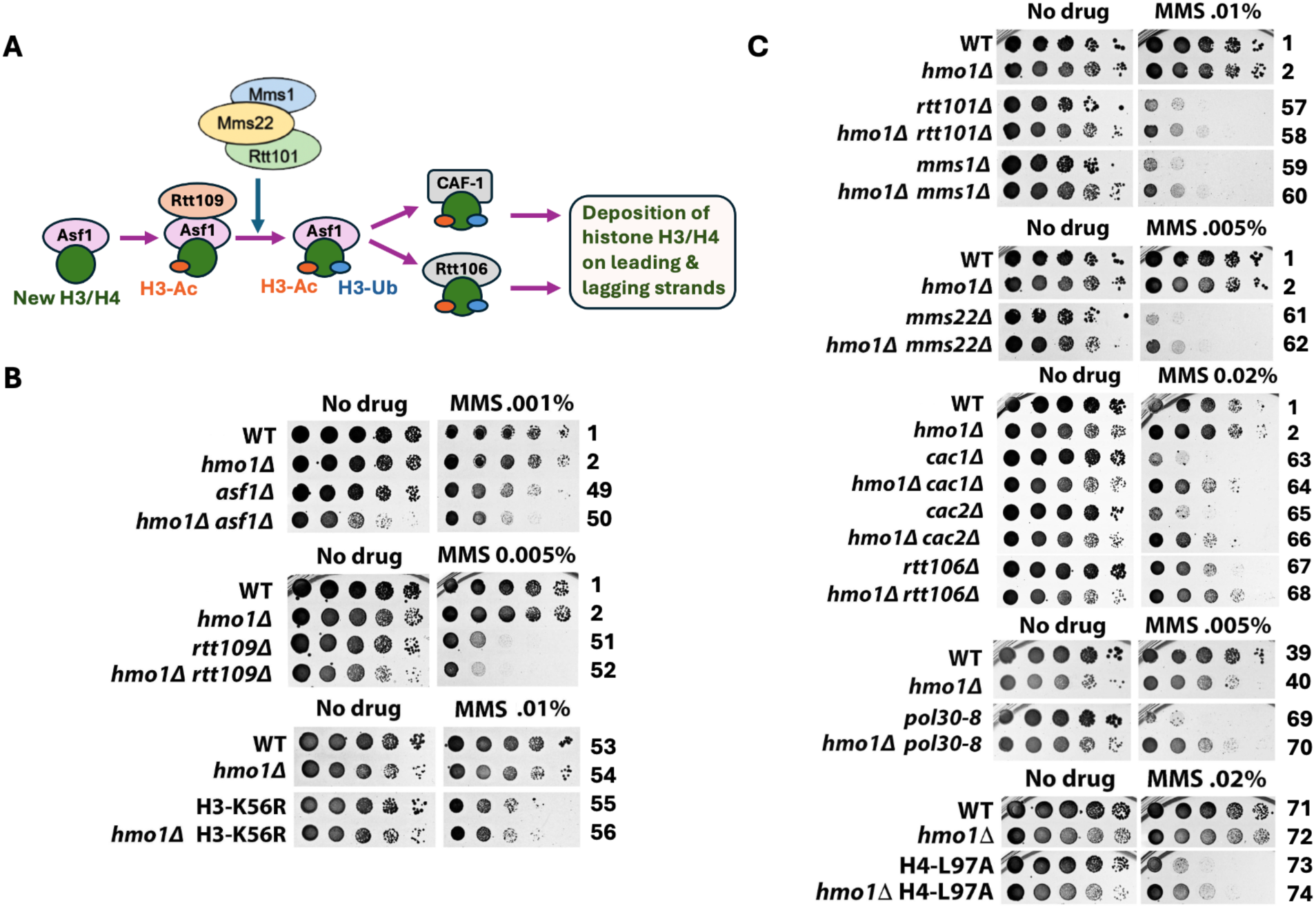
Hmo1 can contribute to DDT by modulating new histone deposition on nascent strand during DNA replication. (A). Mechanism replication-coupled deposition of newly synthesized histones H3/H4 on DNA. H3-Ac, histone H3-K56 acetylation. H3-Ub, H3-K121,122,125 ubiquitination. See the text for descriptions. (B) and (C). Growth phenotypes of indicated strains in the presence or absence of MMS.

To examine whether Hmo1 impacts DDT through modulating replication-coupled chromatin assembly, we examined the interplay of Hmo1 with components of parental histone transfer and new histone deposition pathways in cellular tolerance to MMS. We showed that *hmo1Δ* partially suppressed MMS-sensitivity of *pol32Δ* mutant (Fig. 4B). Pol32 interacts with PCNA and disrupting this interaction by PCNA mutation *pol30-79* also sensitizes cells to MMS (Eissenberg et al. 1997; Serra-Cardona et al. 2024; note PCNA is a homotrimer of Pol30). We found *hmo1Δ* also suppressed MMS-sensitivity of *pol30-79* mutant (Fig. 4B). These results placed Hmo1 upstream of Pol32 in PCNA-Pol δ mediated pathway of parental histones H3/H4 transfer to lagging strand. We found *CTF4* to be epistatic to *HMO1* in MMS tolerance (Fig. 4B), which implicates Hmo1 in MCM-Ctf4-Polα mediated parental histone transfer pathway. We noted that *dpb3Δ* has little effect on cellular tolerance to MMS (Fig. 4B), which is consistent with similar findings in prior studies (Dolce et al. 2022). This is despite that *dpb3Δ* disrupts parental histone H3/H4 transfer to leading strand (Yu et al. 2018). This could be because during DNA replication, ssDNA gaps subject to repair by DDT machinery are more likely to form on lagging strand than leading strand (Pham et al. 2023). Deletion of Hmo1 did not significantly affect the survival of *dpb3Δ* mutant in the presence of MMS (Fig. 4B). In addition, we found *MRC1* to be epistatic to *HMO1* in MMS-tolerance (Fig. 4C), suggesting that Hmo1 functions downstream of Mrc1 in regulating parental histone transfer. These results suggest that Hmo1 functions in parental histone recycling, especially lagging strand-specific histone transfer that seems to impact DDT more than leading strand-specific histone transfer.

We observed an additive/synergistic effect of *hmo1Δ* with *asf1Δ* or *rtt109Δ* on unperturbed cell growth (Fig. 5B, “No drug” panel, note *hmo1Δ asf1Δ* or *hmo1Δ rtt109Δ* double mutants grew significantly slower than its respective single mutants), indicating that Hmo1 contributes to cell growth independently of Asf1 and Rtt109. The survival/growth of *asf1Δ* and *rtt109Δ* mutants in the presence of MMS was reduced by *hmo1Δ* to a degree similar to *hmo1Δ-*induced growth reduction in the absence of MMS (Fig. 5B). As such, *hmo1Δ* did not significantly affect MMS-sensitivity of *asf1Δ* or *rtt109Δ* mutant. Likewise, *hmo1Δ* did not affect MMS-sensitivity of the H3-K56R mutant (Fig. 5B). Therefore, *ASF1* and *RTT109* are epistatic to *HMO1* in MMS-tolerance, suggesting that Hmo1 functions in new histone deposition pathway downstream of Asf1/Rtt109-mediated H3-K56 acetylation. To further test this notion, we examined the interplay of Hmo1 with Rtt101/Mms1/Mms22, CAF-1 and Rtt106 that function down stream of Asf1/Rtt109 (Fig. 5A). As shown in Fig. 5C, *hmo1Δ* suppressed MMS-sensitivity of each of *rtt101Δ*, *mms1Δ, mms22Δ, cac1Δ, cac2Δ* and *rtt106Δ* mutants. CAF-1 recruitment to replicating DNA is mediated by Cac1 interaction with PCNA, and disrupting this interaction by *pol30-8* mutation results in MMS-sensitivity (Zhang et al., 2000). We showed that *hmo1Δ* also suppressed MMS-sensitivity of *pol30-8* mutant (Fig. 5C). The histone H4-L97A mutation hinders the deposition of histone H3/H4 dimer to DNA by CAF-1 or Rtt106 (Yu et al. 2011). We found *hmo1Δ* to also suppress MMS-sensitivity of cells bearing the histone H4-L97A mutation (Fig. 5C). These results suggest that Hmo1 can contribute to DTT by functioning downstream of Asf1/Rtt109 and upstream of Rtt101/Mms1/Mms22 in the new histone deposition pathway.

### Hmo1 antagonizes the function of histone H2A variant H2A.Z (Htz1) in DDT

The highly conserved histone H2A variant H2A.Z (aka Htz1 in yeast) plays multiple functions in genome maintenance and function (Martire and Banaszynski 2020; Ferrand et al. 2020). Histone H2A.Z deposition and removal are mediated by chromatin remodelers SWR-C and INO80, respectively (Mizuguchi et al. 2004; Kobor et al. 2004; Papamichos-Chronakis et al. 2011) (Fig. 6A). There is evidence implicating H2A.Z in cellular response to replicative stress (Srivatsan et al. 2018). Yeast Htz1 deposition and retention in chromatin were found to prevent replisome uncoupling at stalled replication forks, avoiding fork collapse (Srivatsan et al. 2018). We examined the interplay of Hmo1 with Htz1 and Swr1 (catalytic subunit of SWR-C) and found that *hmo1Δ* suppressed MMS-sensitivity of both *htz1Δ* and *swr1Δ* mutants (Fig. 6B). A previous survey of genome-wide distribution of Hmo1 found an inverse correlation between Hmo1 association and Htz1 binding (Bermejo et al. 2009), suggesting a competitive/antagonistic relationship between Hmo1 and Htz1 in their functions in the genome. In support of this notion, we found that the *INO80* gene encoding Ino80, the catalytic subunit of INO80 complex, was epistatic to *HMO1* in MMS tolerance (Fig. 6B), implicating Hmo1 in the Htz1 removal pathway mediated by INO80 complex.

**Fig. 6.**
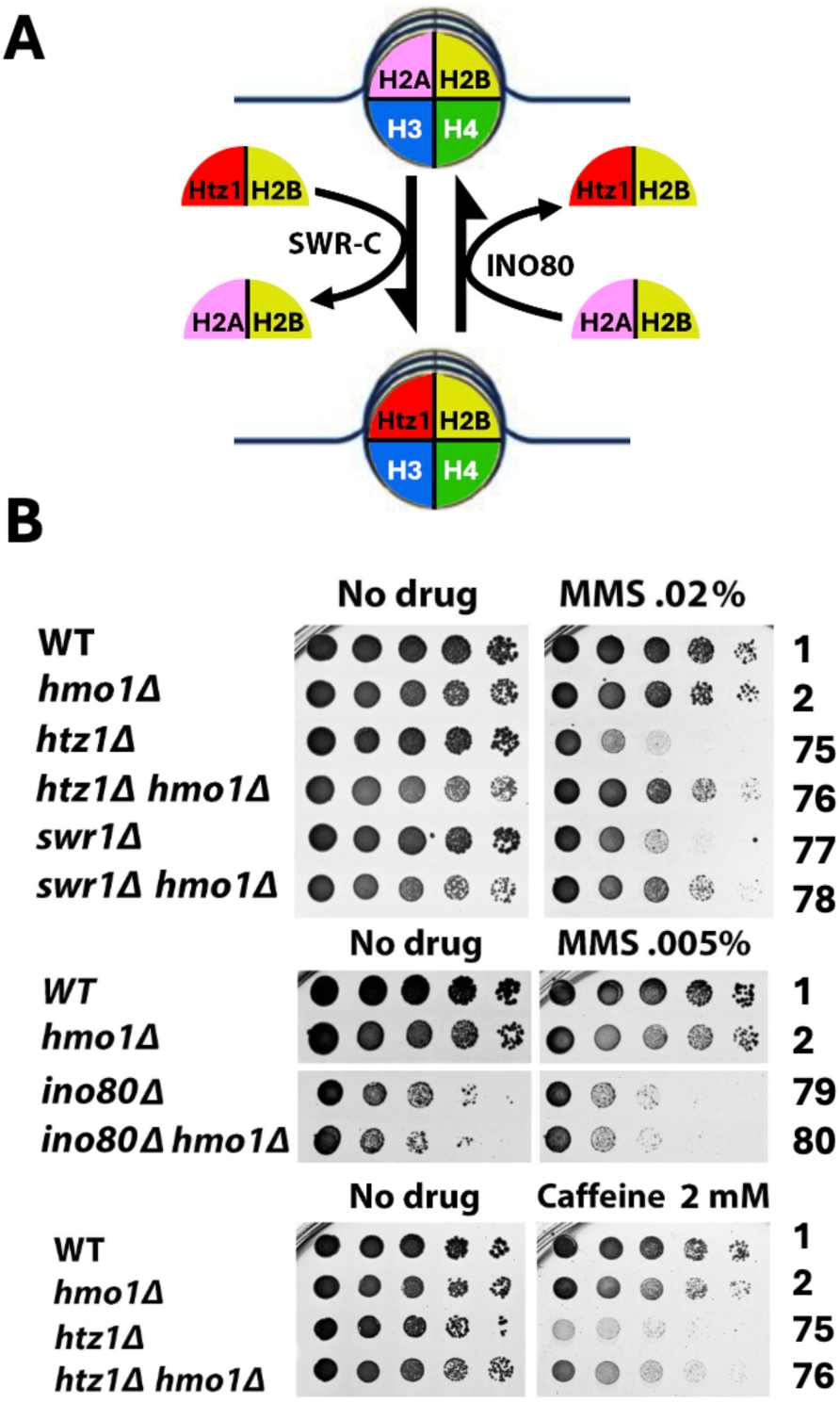
Hmo1 antagonizes the function of histone H2A variant H2A.Z (Htz1) in DDT. (A). Mechanisms of the deposition and removal of H2A.Z (Htz1) from chromatin. See the text for descriptions. (B). Growth phenotypes of indicated strains in the presence or absence of MMS.

Consistent with an antagonistic relationship between Htz1 and Hmo1, they have opposing effects on DNA resection in DSB repair, with Htz1 promoting, and Hmo1 inhibiting, resection (Kalocsay et al. 2015; Panday et al. 2015). The *htz1Δ* mutant is hypersensitive to caffeine that inhibits DNA resection (Mizuguchi et al. 2004; Tsabar et al. 2015), likely because the *htz1Δ* mutation and caffeine together reduced DNA resection to a level too low to allow effective repair of spontaneous DNA damages generated in cell proliferation. We showed that the hypersensitivity of *htz1Δ* mutant to caffeine was partially suppressed by *hmo1Δ* (Fig. 6B), possibility due to an increase in DNA resection as a result of removing the inhibitory effect of Hmo1. It is possible that suppression of MMS-sensitivity of *htz1Δ* mutant by *hmo1Δ* (Fig. 6B) reflects an antagonistic relationship of Htz1 and Hmo1 in impacting DNA resection at the stalled replication fork during DDT.

### Genetic background-dependent differential requirement of CTD of Hmo1 for its functions in DDT

Hmo1 has two HMGB motifs (referred to as domains A and B hereafter) and a basic CTD spanning residues 209-246 (Panday and Grove 2017; Bi 2024) (Fig 7A). Domain A and, to a lesser extent, domain B, bind DNA with a specificity for altered DNA structures including DNA with nicks, gaps, overhangs, or loops, as well as for 4-way junction structures and supercoiled DNA (Albert et al. 2013; Kamau et al. 2004). Hmo1 also bends DNA, which requires its CTD (Bauerle et al, 2006; Xiao et al, 2010). Deletion of CTD does not affect Hmo1 stability or its DNA binding ability (Kamau et al. 2004; Bauerle et al, 2006; Xiao et al, 2010). Chromatin in cells bearing Hmo1 deleted for CTD, referred to as hmo1-AB, was hypersensitive to nuclease digestion, similarly as chromatin in cells deleted for Hmo1, suggesting that CTD is required for the ability of Hmo1 to compact chromatin (Panday and Grove 2016). There is also evidence suggesting that deletion of Hmo1 or its CTD results in a more dynamic chromatin environment at DSBs (Panday et al. 2015). However, whereas Hmo1 deletion results in a salient growth defect in yeast, cells expressing *hmo1-AB* exhibit no growth defect (Bauerle et al, 2006; Xiao et al, 2010). As such, function(s) of Hmo1 required for normal cell proliferation does not reside within the CTD.

**Fig. 7.**
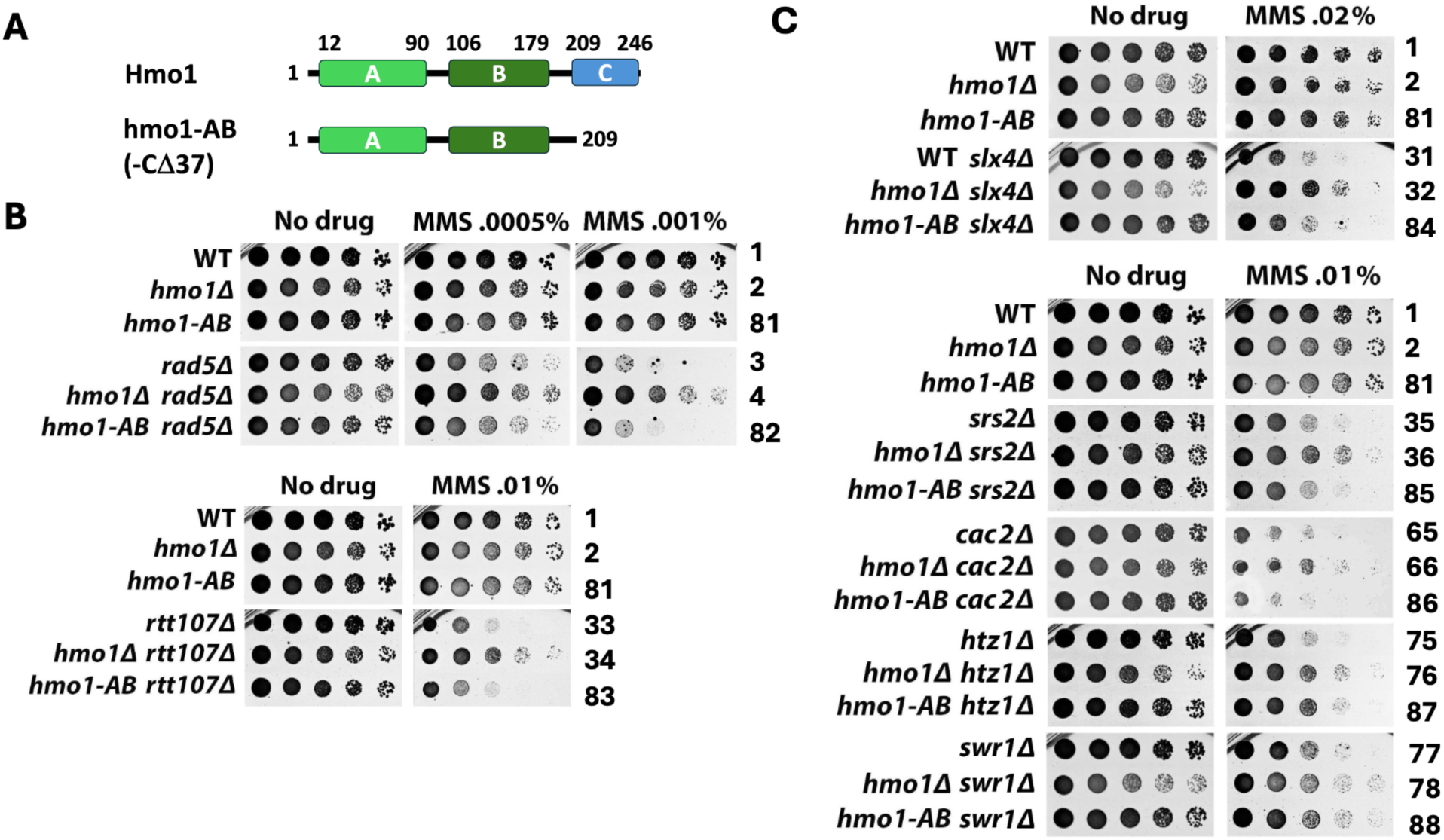
The CTD of Hmo1 contributes to DDT in a genetic background-dependent manner. (A). Domain structure of Hmo1. Top, domains (HMGB motifs) A and B, and the basic CTD of Hmo1. Bottom, the hmo1-AB allele (Hmo1 deleted for the CTD). (B) and (C). Growth phenotypes of indicated strains in the presence or absence of MMS.

To test if CTD of Hmo1 is required for its function in DDT, we examined if the *hmo1-AB* mutation resembles *hmo1Δ* in affecting MMS-tolerance. We first tested if *hmo1-AB* could suppress MMS-sensitivity of *rad5Δ* mutant, similarly as *hmo1Δ*. As shown in Fig. 7B, unlike *hmo1Δ*, *hmo1-AB* did not suppress MMS-sensitivity of *rad5Δ* mutant. We found that *hmo1-AB* also failed to suppress MMS-sensitivity of *rtt107Δ, slx4Δ, srs2Δ* or cac2*Δ* mutant (Fig. 7B and 7C). However, like *hmo1Δ*, *hmo1-AB* suppressed MMS sensitivity of *htz1Δ* and *swr1Δ* mutant (Fig. 7C). These results indicate that CTD of Hmo1 is dispensable for its function in DDT in cells lacking Rad5, Rtt107, Slx4, Srs2 or Cac2, but is necessary for Hmo1 function in cells lacking Htz1 or Swr1. Note in the above experiment, *hmo1-AB* mutation was introduced to the endogenous *HMO1* locus of the host. We were able to reproduce the results demonstrating the ability of *hmo1-AB* to suppress *htz1Δ* but not *rad5Δ* or *rtt107Δ* by expressing the *hmo1-AB* allele carried on a plasmid (Fig. S1).

Our finding that *hmo1-AB* didn’t suppress MMS-sensitivity of *rad5Δ* mutant is in not consistent with an earlier report showing that deletion of C-terminal 64 or 22 residues of Hmo1 (hmo1-Δ64 or hmo1-Δ22) partially suppressed MMS-sensitivity of *rad5Δ* mutant (Gonzalez-Huici et al. 2014). The *hmo1-AB* allele we used here has been characterized in several prior investigations on Hmo1 functions (Xiao et al. 2010, 2011; Panday et al. 2015, 2017). It is deleted for C-terminal 37 residues (from 209 to 246) of Hmo1, more than that in the hmo1-Δ22 allele but less than that in hmo1-Δ64 used by Gonzalez-Huici et al. (Gonzalez-Huici et al. 2014). Therefore, the discrepancy between our result and that of Gonzalez-Huici et al. is unlikely due to the difference in the size of the truncation of the Hmo1 alleles. We noted that the strains we used were derived from BY4741 (Table 1) whereas those used by Gonzalez-Huici et al. were W303 derivatives (Gonzalez-Huici et al. 2014). It is possible that the discrepancy regarding the role of Hmo1-CTD in MMS-tolerance in *rad5Δ* mutant stemmed from the difference in the genetic background between BY4741 and W303 (Sherman 2002).

### Hmo1 promotes DNA damage checkpoint signaling induced by replication stress

Replication stress induces S-phase checkpoint that is initiated by Mec1 kinase and facilitated by checkpoint mediators Rad9 or Mrc1 (Pardo et al. 2017). It has been shown that Rad9 is the major checkpoint mediator in the presence a low dose of MMS (Bacal et al. 2018). Upon activation via Mec1-dependent phosphorylation, Rad9 binds and activates the checkpoint effector Rad53 kinase which then undergoes intermolecular autophosphorylation (Pardo et al. 2017). Rad53 is also phosphorylated by Mec1 (Pardo et al. 2017). Rad9 is recruited to damaged chromatin by Dpb11 and γH2A (histone H2A phosphorylated at S129 which is equivalent to γH2AX in higher eukaryotes) (Pardo et al. 2017) (Fig. 8A). Notably, the DNA repair scaffold Slx4/Rtt107 is also recruited to DNA lesions by Dpb11 and γH2A (Williams et al. 2010; Ohouo et al. 2010; Mejia-Ramirez et al. 2015) (Fig. 8A). The competition between Rad9 and Slx4/Rtt107 for binding γH2A and Dpb11 is thought to yield a dynamic balance between Rad9-merdiated checkpoint signaling and Slx4/Rtt107-promoted DNA recombination repair (Ohouo et al. 2013) (Fig. 8A). Along this line, *rtt107Δ* or *slx4Δ* results in heightened checkpoint signaling reflected by an increase of Rad9 phosphorylation (Rad9-P) and Rad53 phosphorylation (Rad53-P) induced by MMS (Ohouo et al. 2013; Siler et al. 2024). We have recently found that blocking histone H2A-S129 phosphorylation by the *hta-S129\** mutation also increases the levels of Rad9-P and Rad53-P induced by MMS (Siler et al. 2024). This led to the notion that Slx4/Rtt107 outcompetes Rad9 for binding γH2A at DNA lesions induced by MMS, and consequentially, γH2A plays a net negative role in DDC in response to MMS (Siler et al. 2024).

**Fig. 8.**
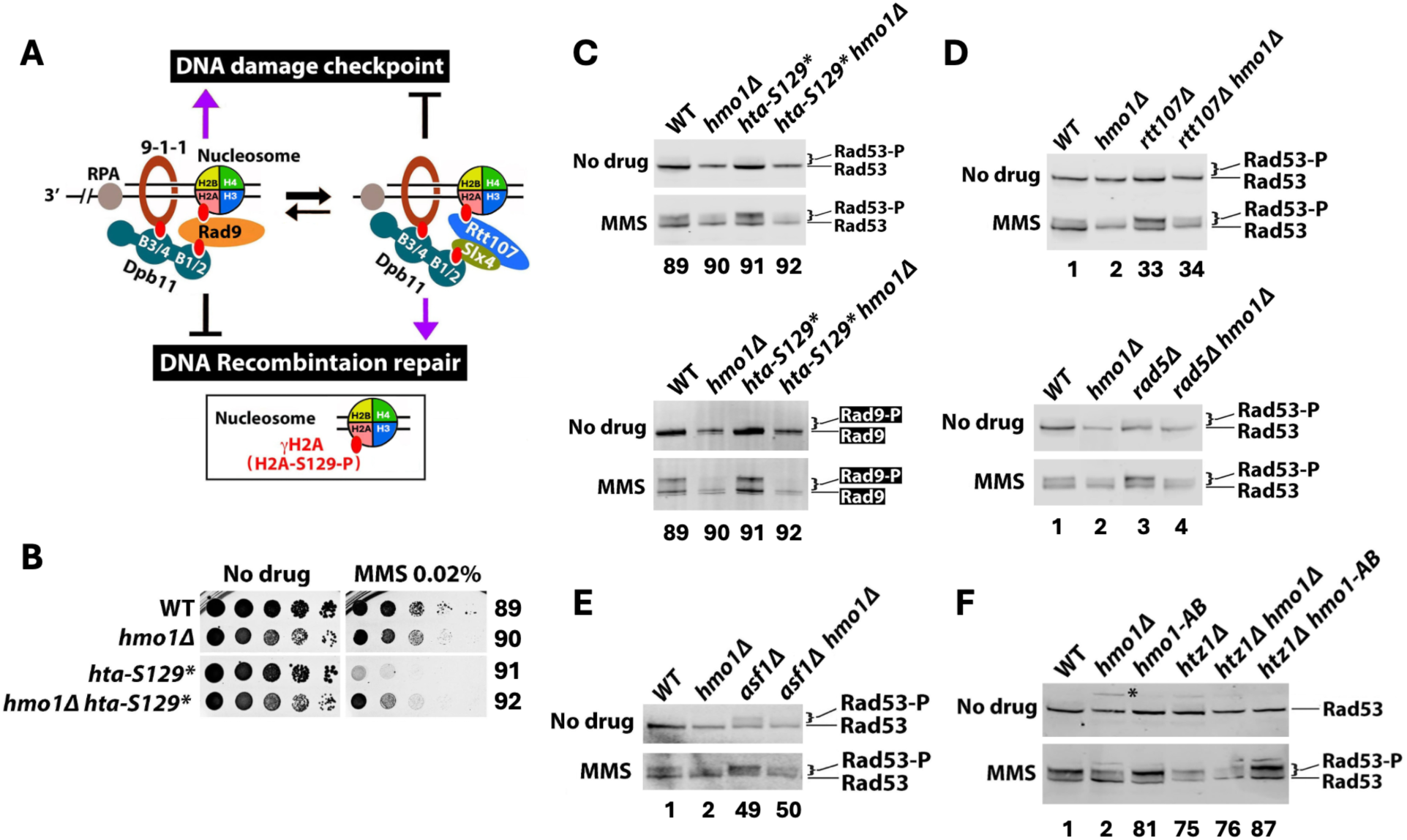
Hmo1 promotes DNA damage checkpoint signaling induced by replication stress. (A) Model for a dynamic balance between DNA damage checkpoint signaling and DNA recombination repair achieved by the competition between Rad9 and Slx4/Rtt107 for binding γH2A and Dpb11 at damaged chromatin. See the text for descriptions. (B) Growth phenotypes of indicated strains in the presence or absence of MMS. (C) to (F). Results from Western blot analyses of Rad53 and Rad9-HA from indicated strains with or without MMS treatment. Log phase cells were treated with (MMS) or without (No drug) MMS at 0.02% for 90 minutes. Protein extract form each cell culture was subjected to SDS-PAGE and Western blotting followed by detection of Rad53 and Rad9-HA with Rad53 and HA antibodies, respectively. Rad53-P, phosphorylated Rad53 protein; Rad9-P, phosphorylated Rad9-HA protein.

The *slx4Δ*, *rtt107Δ*, and *hta-S129\** mutants were all hypersensitive to MMS (Fig. 3 and Fig. 8B), which could be caused by heightened checkpoint and/or reduced recombination repair in them. We found that the MMS sensitivity of *htaS-129\** mutant, similarly as that of *slx4Δ* or *rtt107Δ* mutant, was suppressed by *hmo1Δ* (Fig. 3 and Fig. 8B). We reason that if excessive checkpoint activation in these mutants caused MMS-sensitivity, then suppression of MMS-sensitivity by *hmo1Δ* should be accompanied by a reduction in checkpoint signaling. Consistent with this notion, we found that the level of MMS-induced Rad53-P (relative to unphosphorylated Rad53) in *rtt107Δ hmo1Δ* or *hta-S129* hmo1Δ* double mutant was significantly lower than that in *rtt107Δ* or *hta-S129\** single mutant, respectively (Fig. 8C and 8D). Moreover, the level of MMS-induced Rad9-P in *hta-S129* hmo1Δ* double mutants was markedly lower than that in *hta-S129\** single mutant (Fig. 8C). As such, *rtt107Δ* or *hta-S129\** was no longer able to induce hyperactivation of DDC signaling in the absence of Hmo1. Notably, we also found that MMS-induced Rad53-P and Rad9-P in *hmo1Δ* mutant was markedly lower than that in *HMO1* wildtype cells (Fig. 8C), indicating that Hmo1 has a general role of promoting Rad9-mediated checkpoint activation in response to MMS.

Taken together, the above results are consistent with the notion that excessive DDC activation induced by MMS reduces the survivability of the host, which is subject to suppression by alleviating DDC (e.g., via deleting Hmo1). The *rad5Δ* and *asf1Δ* mutations also caused enhanced DDC as reflected by increased MMS-induced Rad53-P (Fig. 8D and 8E). We found that *hmo1Δ* reduced MMS-induced Rad53-P in *rad5Δ* mutant (Fig. 8D) and suppressed its MMS sensitivity (Fig. 2). However, *hmo1Δ* reduced MMS-induced Rad53-P in *asf1Δ* mutant (Fig. 8E) without suppressing its MMS sensitivity (Fig. 5B). Therefore, whether elevated DDC causes MMS-sensitivity may be dependent on the genetic background of the host.

To gain insight into how Hmo1 promotes DDC, we tested if CTD of Hmo1 is required for this function. We found that unlike *hmo1Δ,* the *hmo1-AB* mutation did not reduce the level Rad53-P induced by MMS (Fig. 8F). Therefore, the ability of Hmo1 to promote DDC does not depend on its CTD and by inference DNA bending activity. While *hmo1-AB* suppressed MMS-sensitivity of *htz1Δ* mutant similarly as *hmo1Δ* (Fig. 7C), it did not reduce Rad53-P in *htz1Δ* mutant as did *hmo1Δ* (Fig. 8F). This indicates that Hmo1’s function in DDT in *htz1Δ* mutant is independent of its ability to promote checkpoint signaling.

### Hmo1 regulates DDT independently of its impact on DNA supercoiling as a proxy of chromatin structure

Hmo1 has been shown to alter nucleosome conformation *in vitro* and promote chromatin compaction *in vivo* (Lu et al. 1996; McCauley et al. 2019; Malinina et al. 2022; reviewed in Bi 2024). Since nucleosome formation introduces negative supercoiling into DNA, the level of DNA negative supercoiling can be used as an indicator of chromatin structure (Simpson et al. 1985). We found that *hmo1Δ* reduced the negative supercoiling of 2-micron plasmid (2μ) in yeast by a linking number difference (ΔLk) of 1 (Fig. 9), suggesting that Hmo1 promotes nucleosome/chromatin formation and/or maintenance. Whether Hmo1 regulates DDT via impacting chromatin has not been investigated. To address this question, we examined if the interplay of Hmo1 with DDT factors, especially those known to modulate the structure and/or replication of chromatin, is linked to the ability of Hmo1 to promote negative DNA supercoiling as a proxy of chromatin structure.

**Fig. 9.**
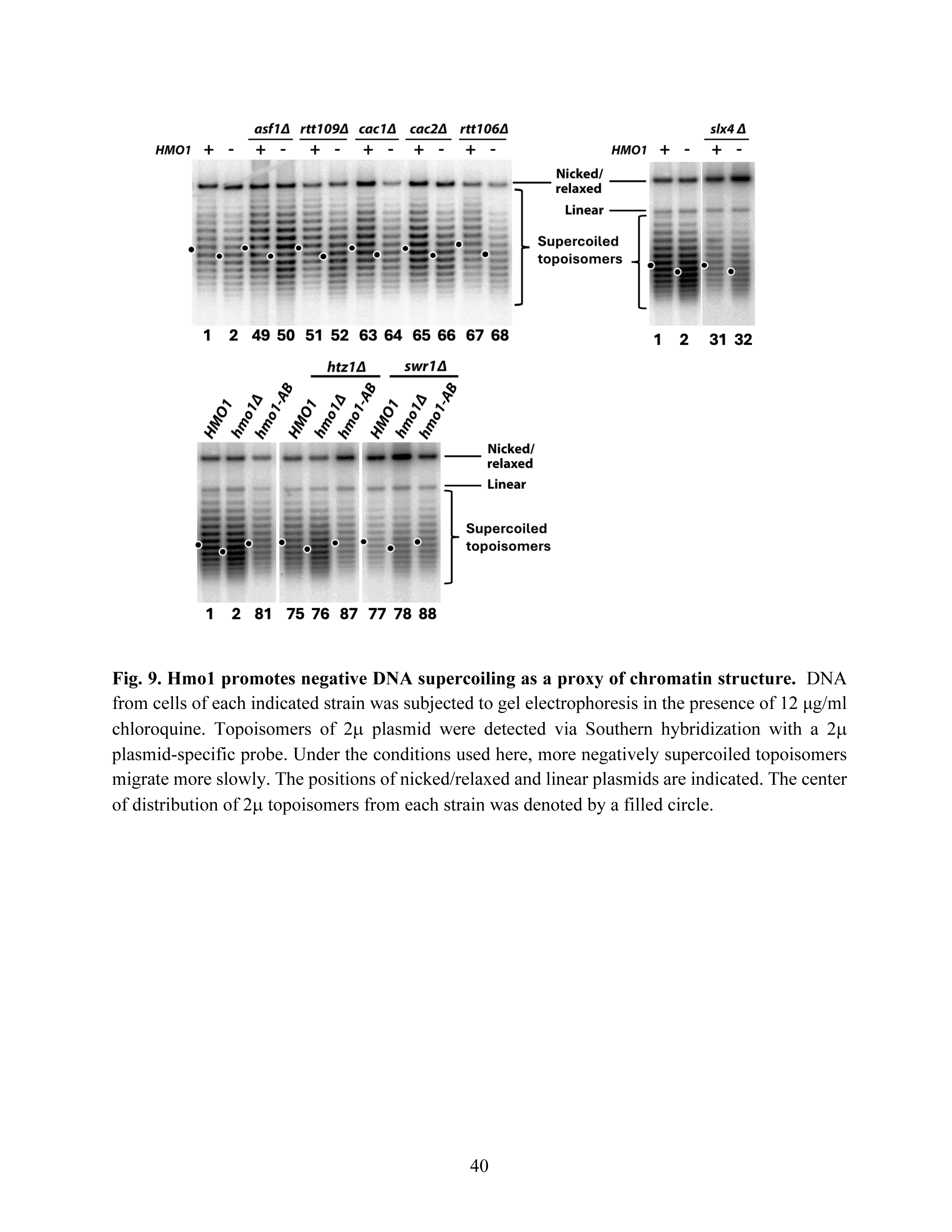
Hmo1 promotes negative DNA supercoiling as a proxy of chromatin structure. DNA from cells of each indicated strain was subjected to gel electrophoresis in the presence of 12 μg/ml chloroquine. Topoisomers of 2μ plasmid were detected via Southern hybridization with a 2μ plasmid-specific probe. Under the conditions used here, more negatively supercoiled topoisomers migrate more slowly. The positions of nicked/relaxed and linear plasmids are indicated. The center of distribution of 2μ topoisomers from each strain was denoted by a filled circle.

We found that none of the *asf1Δ, rtt109Δ, cac1Δ, cac2Δ, rtt106Δ, slx4Δ, htz1Δ,* and *swr1Δ* mutations affected 2μ supercoiling (Fig. 9). As such, MMS-sensitivity caused by these mutations are not due to a change in general chromatin structure. Deletion of *HMO1* from any of these mutants reduced 2μ supercoiling by a ΔLk of 1, similarly as it did in the otherwise wildtype strain (Fig. 9). Note that *hmo1Δ* suppressed MMS-sensitivity of *cac1Δ, cac2Δ, rtt106Δ, slx4Δ,* and *htz1Δ* mutants (Fig. 3, 5C, and 6B) but did not affect MMS-sensitivity of *asf1Δ* or *rtt109Δ* mutant (Fig. 5A). These results suggest that how Hmo1 functions in DDT is not linked to its role in promoting DNA supercoiling as a proxy of chromatin structure.

We wondered if DNA bending mediated by Hmo1-CTD is required for the function of Hmo1 in promoting DNA supercoiling. To address this question, we tested the effect of *hmo1-AB* mutation on 2μ supercoiling. We found that, unlike *hmo1Δ*, *hmo1-AB* mutation did not affect 2μ supercoiling (Fig. 9), indicating that CTD of Hmo1 is dispensable for its ability to promote DNA supercoiling. We also showed that *hmo1-AB* mutation did not affect 2μ supercoiling in *htz1Δ* or *swr1Δ* mutant (Fig. 9). That both *hmo1Δ* and *hmo1-AB* mutations suppressed MMS-sensitivity of *htz1Δ* and *swr1Δ* mutants (Fig. 7C) but only *hmo1Δ* reduced DNA supercoiling in these mutants (Fig. 9) supports the notion Hmo1 can regulate DDT independently of its impact on DNA supercoiling as a proxy of chromatin structure.

## Discussion

### Hmo1 may direct persistent SCJs to Mus81/Mms4 nuclease-mediated resolution pathway

We found evidence implicating Hmo1 in SCJ resolution pathway mediated by the Slx4-Mus81/Mms4 axis instead of the parallel SCJ dissolution pathway mediated by the Smc5/6-STR axis (Fig. 1). The suppressive effect of *hmo1Δ* on MMS-sensitivity of *mus81Δ*, *mms4Δ*, and *slx4Δ* mutants can be explained by assuming that Hmo1 channels SCJ induced by MMS to Mus81/Mms4-dependent resolution pathway. It is noteworthy that both STR dissolvase and Mus81/mms4 resolvase are cell cycle regulated. STR is subject to S-phase specific Smc5/6-mediated SUMOylation that is required for efficient function of STR (Bermúdez-López et al. 2016), whereas Mus81/mms4 activation depends on the phosphorylation of Mms4 and the formation of a multiprotein complex containing Mus81/Mms4 and Slx4/Dpb11 in G2/M phase (Gallo-Fernández et al. 2012; Szakal and Branzei 2013; Gritenaite et al. 2014). As such, it is believed that STR acts in S phase to dissolve SCJs thus preventing unscheduled recombination during DNA replication that may lead to DNA rearrangement, while Mus81/mms4 is employed in G2/M phase to resolve SCJs that escape STR action (Szakal and Branzei 2013). We envision that Hmo1 may act upstream of Mus81/Mms4 in SCJ resolution pathway by promoting the formation and/or stability of dHJs. Hmo1 may aid in the conversion of SCJ to dHJs by facilitating the reannealing of parental strands in SCJ (Fig. 1). Alternatively, or in addition, Hmo1 may bind and stabilize preformed dHJs. In any case, when Mus81/Mms4 is absent, unresolved dHJs accumulate and become toxic, which can be alleviated by Hmo1 deletion that reduces the abundance of dHJs destinated to be processed by Mus81/Mms4.

### Hmo1 may regulate DDT by impacting replication-coupled chromatin assembly

DDT is intimately intertwined with DNA replication and chromatin packaging. As a sequence non-specific DNA binding protein that preferentially binds DNA with altered conformations, Hmo1 may regulate the formation, stability, and/or processing of special DNA structures generated during ordinary DNA replication or replication stress. These structures include ssDNA gaps, D-loops, SCJs, and dHJs. In addition, since Hmo1 can bind and alter nucleosome structure (McCauley et al. 2019; Malinina et al. 2022), it may modulate nucleosome assembly that is tightly coupled with ongoing DNA replication (Zhang et al. 2020). It is reasonable to posit that the ability of Hmo1 to bind altered DNA structures and that to modulate nucleosome assembly/structure mutually influence each other. The interplay of Hmo1 with replication-coupled chromatin assembly machinery in MMS tolerance we observed (Fig. 4 and Fig. 5) suggests that Hmo1 can modulate DDT by functioning in both parental histone transfer and new histone deposition in the wake of replication fork. Hmo1 likely acts upstream of Pol32 in Pol δ-mediated parental histone transfer to lagging strand and upstream of CAF-1 and Rtt106 in new histone deposition pathway to ensure proper chromatin assembly during DNA replication. In doing so, Hmo1 likely promotes the creation of a state of nascent chromatin at stalled replication fork that renders DNA lesions or ssDNA gaps more accessible to TS factors such as Rad5 and Mph1 than the TLS machinery, thereby channeling DNA lesions to TS pathway (Fig. 1). Similarly, Hmo1 may also aids in the establishment of a chromatin environment at SCJ that is conducive to its conversion to dHJs destinated to resolution by Mus81/Mms4 nuclease (Fig. 1).

### Hmo1 counteracts the function of histone H2A variant H2A.Z (Htz1) in DDT

Our finding that *hmo1Δ* suppresses hypersensitivity of *htz1Δ* mutant to caffeine, an inhibitor of DNA resection (Fig. 6), is in line with the antagonistic relationship of Htz1 and Hmo1 on DNA resection involved in DSB repair (Kalocsay et al. 2009; Panday et al. 2015). Htz1 promotes DSB repair likely because it makes nucleosome/chromatin structure more accessible (Kalocsay et al. 2009). On the other hand, Hmo1 inhibits DSB end resection, possibly by stabilizing chromatin (Panday et al. 2015). Based on the inverse correlation between Htz1 and Hmo1 in genome association (Bermejo et al. 2009), we propose that Htz1 counteracts Hmo1 binding to DNA/chromatin, thereby alleviating its inhibition of DSB repair. Consistent with this notion, Htz1 is transiently recruited to DSBs and Hmo1 is evicted during DSB repair (Kalocsay et al. 2009; Panday et al. 2015). There is evidence implicating DNA resection in the formation/expansion of ssDNA gaps at stalled replication forks (Ball et al. 2014; Hale et al. 2023; García-Rodríguez et al. 2024). It is tempting to assume that, similarly as DNA resection in DSB repair, DNA resection involved in DDT is also subject to positive regulation by Htz1 and negative regulation by Hmo1. This hypothesis can explain the suppression of MMS-sensitivity of *htz1Δ* mutant by *hmo1Δ* (Fig. 6).

### Contribution of the CTD of Hmo1 to DDT is dependent on the genetic background of the host

The molecular mechanisms underlying Hmo1’s functions in DDT have yet to be elucidated. Hmo1 has both HMGB-like characteristics and linker histone-like characteristics (Panday and Grove 2017; Bi 2024). It’s not clear how any of these characteristics may contribute to Hmo1 function in DDT. The lysine rich basic CTD of Hmo1 resembles that of canonical linker histones and is believed to be the structural basis of Hmo1’s linker histone-like characteristics (Panday and Grove 2017; Bi 2024). CTD of Hmo1 is required for its DNA bending activity but is dispensable for its DNA-binding activity (Bauerle et al, 2006; Kamau et al. 2004; Xiao et al. 2010; Panday et al. 2015). CTD and by inference DNA bending is required for Hmo1’s abilities to make yeast genome resistant to nuclease digestion possibly due to chromatin compaction (Panday and Grove 2016) and to stabilize chromatin (promoting a less dynamic chromatin environment) (Panday et al. 2015). However, we found that CTD of Hmo1 is not required for its function in promoting negative supercoiling of DNA (Fig. 9) which is an indicator of the formation of nucleosomes/primary chromatin (Simpson et al. 1985). Therefore, Hmo1 seems to contribute to the formation of primary chromatin and chromatin compaction via distinct CTD-independent and CTD-dependent mechanisms, respectively. CTD of Hmo1 has recently been found to be necessary for its interaction with chromatin remodeling complexes (Amigo et al. 2022), which raises the possibility that Hmo1 promotes chromatin compaction by recruiting chromatin remodeling complexes.

That *hmo1-AB* phenocopied *hmo1Δ* in suppressing MMS-sensitivity of *htz1Δ* or *swr1Δ* mutant (Fig. 7C) pointed to a role of the CTD of Hmo1 in DDT. However, unlike *hmo1Δ*, *hmo1-AB* failed to suppress MMS-sensitivity of *rad5Δ, rtt107Δ, slx4Δ, srs2Δ* or cac2*Δ* mutant (Fig. 7B and 7C), demonstrating that CTD is dispensable for Hmo1 function in these mutants. These results indicate that Hmo1 may function in DDT via CTD-dependent or CTD-independent mechanisms depending on the genetic background of the host. The *htz1Δ* mutant is hypersensitive to caffeine that inhibits DNA resection (Mizuguchi et al. 2004; Tsabar et al. 2015), which is suppressed by *hmo1Δ* (Fig. 6B). On the other hand, we found that none of the *rad5Δ, rtt107Δ, slx4Δ,* and *srs2Δ* mutants was sensitive to caffeine (Fig. S2). It is possible the unique circumstance of *htz1Δ* mutant (e.g., reduced DNA resection capability) enables an CTD-dependent function of Hmo1 (e.g., inhibition of DNA resection) in DDT to take place or become manifest. This CTD-dependent Hmo1 function may not take place or be required for DDT in the *rad5Δ, rtt107Δ, slx4Δ,* or *srs2Δ* mutant (in which DNA resection is not reduced as in *htz1Δ* mutant). How and under what circumstances the CTD of Hmo1 contributes to its function in DDT await further investigations.

### Hmo1 promotes DNA damage checkpoint signaling

Our finding that *hmo1Δ* reduced MMS-induced Rad53-P and Rad9-P in yeast (Fig. 8C) demonstrated a role of Hmo1 in promoting DNA damage checkpoint signaling in response to replication stress. Hmo1 may help create a local chromatin environment that is favorable for the functions of DDC kinase Mec1 and/or mediator Rad9. Mec1 is recruited to ssDNA by ssDNA-binding protein RPA (Pardo et al., 2017). As RPA-bound sites in the genome overlap those of Hmo1-associated sites in yeast genome upon MMS treatment (Gonzalez-Huici et al. 2014), it is possible that Hmo1 contributes to the stability of RPA-covered ssDNA regions at stalled replication forks, thereby facilitating Mec1 recruitment. Rad9 is recruited to damaged chromatin in part by associating with γH2A (Pardo et al. 2017). Hmo1 may aids in Rad9 recruitment by altering the structure of γH2A-bearing nucleosomes in a manner that increases the affinity of γH2A for Rad9, instead of its competitive inhibitor Rtt107 (Fig. 8A). Notably, we found that the CTD of Hmo1 and by inference its DNA bending activity is not required for its role in promoting of DDC in response to MMS (Fig. 8F). We found that the function of Hmo1 in DDT in *htz1Δ* mutant is independent on the ability of Hmo1 to promote DDC signaling (Fig. 8F). A decrease in DDC signaling caused by *hmo1Δ* is correlated with an improvement of MMS-tolerance in mutants such as *rtt107Δ* but not in the *asf1Δ* mutant (Fig. 8D and 8E). Therefore, Hmo1-mediated promotion of DDC signaling may be irrelevant to DDT or may be involved in DDT in a genetic background-dependent manner.

## Acknowledgments

We thank the Department of Biology at University of Rochester for support. We thank Drs Xiaolan Zhao (Memorial Sloan Kettering Cancer Center), Stephen Kron (University of Chicago), Dr. Peter Burgers (Washington University), and Anne Grove (Louisiana State University) for their gifts of yeast strains and plasmids.

---

**Fig. S1.**
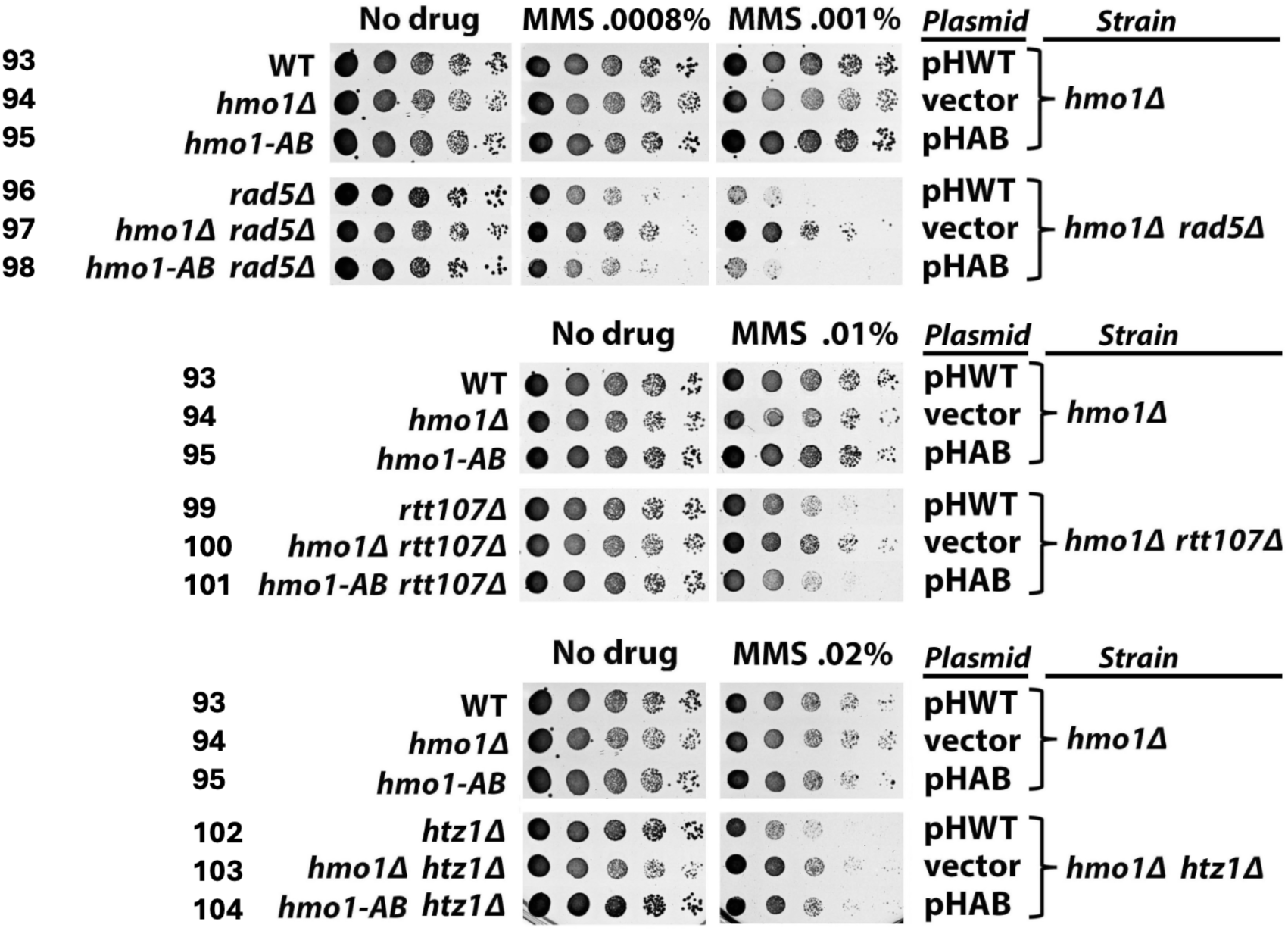
Examination of the role of CTD of Hmo1 in DDT using strains carrying plasmid borne *HMO1* alleles. Shown are growth phenotypes of indicated strains in the presence or absence of MMS.

**Fig. S2.**
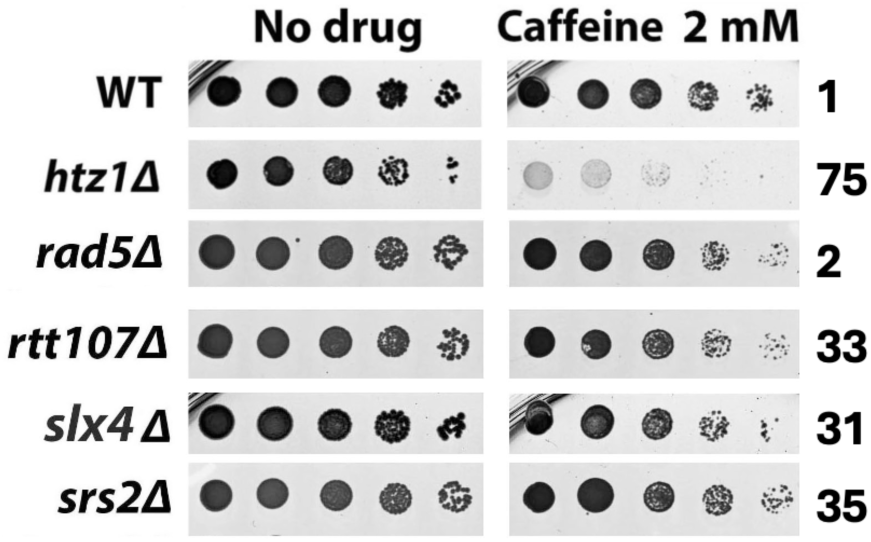
Hmo1, but not Rad5, Rtt107, Slx4, or Srs2 is required for yest tolerance to caffeine. Shown are growth phenotypes of indicated strains on medium with or without caffeine.

## Notes

### Competing Interest Statement

The authors have declared no competing interest.

## References

Achar YJ, Adhil M, Choudhary R, Gilbert N, Foiani M. Negative supercoil at gene boundaries modulates gene topology. Nature 2020; 577: 701–705 [PMID: 31969709 DOI: 10.1038/s41586-020-1934-4]

Albert B, Colleran C, Léger-Silvestre I, Berger AB, Dez C, Normand C, Perez-Fernandez J, McStay B, Gadal O. Structure-function analysis of Hmo1 unveils an ancestral organization of HMG-Box factors involved in ribosomal DNA transcription from yeast to human. Nucleic Acids Res 2013; 41: 10135–10149 [PMID: 24021628 DOI: 10.1093/nar/gkt770]

Amigo R, Farkas C, Gidi C, Hepp MI, Cartes N, Tarifeño E, Workman JL, Gutiérrez JL. The linker histone Hho1 modulates the activity of ATP-dependent chromatin remodeling complexes. Biochim Biophys Acta Gene Regul Mech 2022; 1865: 194781 [PMID: 34963628 DOI: 10.1016/j.bbagrm.2021.194781]

Bacal J, Moriel-Carretero M, Pardo B, Barthe A, Sharma S, Chabes A, Lengronne A, Pasero P. Mrc1 and Rad9 cooperate to regulate initiation and elongation of DNA replication in response to DNA damage. EMBO J. 2018 Nov 2;37(21):e99319. doi: 10.15252/embj.201899319. Epub 2018 Aug 29.PMID: 30158111

Ball LG, Hanna MD, Lambrecht AD, Mitchell BA, Ziola B, Cobb JA, Xiao W. The Mre11-Rad50-Xrs2 complex is required for yeast DNA post replication repair. PLoS One. 2014 Oct 24;9(10):e109292. doi: 10.1371/journal.pone.0109292. eCollection 2014.PMID: 25343618

Bauerle KT, Kamau E, Grove A. Interactions between N- and C-terminal domains of the Saccharomyces cerevisiae high-mobility group protein HMO1 are required for DNA bending. Biochemistry 2006; 45: 3635–3645 [PMID: 16533046 DOI: 10.1021/bi0522798]

Bellelli R, Borel V, Logan C, Svendsen J, Cox DE, Nye E, Metcalfe K, O’Connell SM, Stamp G, Flynn HR, Snijders AP, Lassailly F, Jackson A, Boulton SJ. Polε Instability Drives Replication Stress, Abnormal Development, and Tumorigenesis. Mol Cell. 2018 May 17;70(4):707–721.e7. doi: 10.1016/j.molcel.2018.04.008. Epub 2018 May 10.PMID: 29754823

Bermejo R, Capra T, Gonzalez-Huici V, Fachinetti D, Cocito A, Natoli G, Katou Y, Mori H, Kurokawa K, Shirahige K, Foiani M. Genome-organizing factors Top2 and Hmo1 prevent chromosome fragility at sites of S phase transcription. Cell 2009; 138: 870–884 [PMID: 19737516 DOI: 10.1016/j.cell.2009.06.022]

Bermúdez-López M, Villoria MT, Esteras M, Jarmuz A, Torres-Rosell J, Clemente-Blanco A, Aragon L. Sgs1’s roles in DNA end resection, HJ dissolution, and crossover suppression require a two-step SUMO regulation dependent on Smc5/6. Genes Dev. 2016 Jun 1;30(11):1339–56. doi: 10.1101/gad.278275.116.PMID: 27298337

Bi X. Hmo1: A versatile member of the high mobility group box family of chromosomal architecture proteins World J Biol Chem. 2024 Aug 12;15(1):97938. doi: 10.4331/wjbc.v15.i1.97938.PMID: 39156122

Bi X. Mechanism of DNA damage tolerance. World J Biol Chem 2015; 6: 48–56 [PMID: 26322163 DOI: 10.4331/wjbc.v6.i3.48]

Bizard AH, Hickson ID. The dissolution of double Holliday junctions. Cold Spring Harb Perspect Biol. 2014 Jul 1;6(7):a016477. doi: 10.1101/cshperspect.a016477.PMID: 24984776

Blastyák A, Pintér L, Unk I, Prakash L, Prakash S, Haracska L. Yeast Rad5 protein required for postreplication repair has a DNA helicase activity specific for replication fork regression Mol Cell. 2007 Oct 12;28(1):167–75. doi: 10.1016/j.molcel.2007.07.030.PMID: 17936713

Bonner JN, Choi K, Xue X, Torres NP, Szakal B, Wei L, Wan B, Arter M, Matos J, Sung P, Brown GW, Branzei D, Zhao X. Smc5/6 Mediated Sumoylation of the Sgs1-Top3-Rmi1 Complex Promotes Removal of Recombination Intermediates. Cell Rep. 2016a Jul 12;16(2):368–378. doi: 10.1016/j.celrep.2016.06.015. Epub 2016 Jun 30.PMID: 27373152

Bonner JN, Zhao X. Replication-Associated Recombinational Repair: Lessons from Budding Yeast. Genes (Basel). 2016b Aug 17;7(8):48. doi: 10.3390/genes7080048.PMID: 27548223

Branzei D, Psakhye I. DNA damage tolerance. Curr Opin Cell Biol. 2016; 40:137–44. doi: 10.1016/j.ceb.2016.03.015.

Byun TS, Pacek M, Yee MC, Walter JC, Cimprich KA. Functional uncoupling of MCM helicase and DNA polymerase activities activates the ATR-dependent checkpoint. Genes Dev. 2005; 19: 1040– 1052. PMID: 15833913

Charlton SJ, Flury V, Kanoh Y, Genzor AV, Kollenstart L, Ao W, Brøgger P, Weisser MB, Adamus M, Alcaraz N, Delvaux de Fenffe CM, Mattiroli F, Montoya G, Masai H, Groth A, Thon G. The fork protection complex promotes parental histone recycling and epigenetic memory. Cell. 2024 Sep 5;187(18):5029–5047.e21. doi: 10.1016/j.cell.2024.07.017. Epub 2024 Aug 1.PMID: 39094569

Chavdarova M, Marini V, Sisakova A, Sedlackova H, Vigasova D, Brill SJ, Lisby M, Krejci L. Srs2 promotes Mus81-Mms4-mediated resolution of recombination intermediates. Nucleic Acids Res. 2015 Apr 20;43(7):3626–42. doi: 10.1093/nar/gkv198. Epub 2015 Mar 12.PMID: 25765656

Chen YH, Choi K, Szakal B, Arenz J, Duan X, Ye H, Branzei D, Zhao X. Interplay between the Smc5/6 complex and the Mph1 helicase in recombinational repair. Proc Natl Acad Sci U S A. 2009 Dec 15;106(50):21252–7. doi: 10.1073/pnas.0908258106. Epub 2009 Dec 7.PMID: 19995966

Collins SR, Miller KM, Maas NL, Roguev A, Fillingham J, Chu CS, Schuldiner M, Gebbia M, Recht J, Shales M, Ding H, Xu H, Han J, Ingvarsdottir K, Cheng B, Andrews B, Boone C, Berger SL, Hieter P, Zhang Z, Brown GW, Ingles CJ, Emili A, Allis CD, Toczyski DP, Weissman JS, Greenblatt JF, Krogan NJ. Functional dissection of protein complexesinvolved in yeast chromosome biology using a genetic interaction map. Nature. 2007 Apr 12;446(7137):806-10. doi: 10.1038/nature05649. Epub 2007 Feb 21.PMID: 17314980

Crickard JB, Moevus CJ, Kwon Y, Sung P, Greene EC. Rad54 Drives ATP Hydrolysis Dependent DNA Sequence Alignment during Homologous Recombination. Cell. 2020 Jun 11;181(6):1380–1394.e18. doi: 10.1016/j.cell.2020.04.056. Epub 2020 Jun 4.PMID: 32502392

Dolce V, Dusi S, Giannattasio M, Joseph CR, Fumasoni M, Branzei D. Parental histone deposition on the replicated strands promotes error-free DNA damage tolerance and regulates drug resistance. Genes Dev. 2022 Feb 1;36(3-4):167–179. doi: 10.1101/gad.349207.121. Epub 2022 Feb 3.PMID: 35115379

Eissenberg JC, Ayyagari R, Gomes XV, Burgers PM. Mutations in yeast proliferating cell nuclear antigen define distinct sites for interaction with DNA polymerase delta and DNA polymerase epsilon. Mol Cell Biol. 1997 Nov;17(11):6367–78. doi: 10.1128/MCB.17.11.6367.PMID: 9343398

Ferrand J, Rondinelli B, Polo SE. Histone Variants: Guardians of Genome Integrity. Cells. 2020 Nov 5;9(11):2424. doi: 10.3390/cells9112424.PMID: 33167489

Gaines WA, Godin SK, Kabbinavar FF, Rao T, VanDemark AP, Sung P, Bernstein KA. Promotion of presynaptic filament assembly by the ensemble of S. cerevisiae Rad51 paralogues with Rad52. Nat Commun. 2015 Jul 28;6:7834. doi: 10.1038/ncomms8834.PMID: 26215801

Gallo-Fernández M, Saugar I, Ortiz-Bazán MÁ, Vázquez MV, Tercero JA. Cell cycle-dependent regulation of the nuclease activity of Mus81-Eme1/Mms4. Nucleic Acids Res. 2012 Sep 1;40(17):8325–35. doi: 10.1093/nar/gks599. Epub 2012 Jun 22.PMID: 22730299

Gan H, Serra-Cardona A, Hua X, Zhou H, Labib K, Yu C, Zhang Z. The Mcm2-Ctf4-Polα Axis Facilitates Parental Histone H3-H4 Transfer to Lagging Strands. Mol Cell. 2018 Oct 4;72(1):140–151.e3. doi: 10.1016/j.molcel.2018.09.001. Epub 2018 Sep 20.PMID: 30244834

García-Rodríguez N, Domínguez-García I, Domínguez-Pérez MDC, Huertas P. EXO1 and DNA2-mediated ssDNA gap expansion is essential for ATR activation and to maintain viability in BRCA1-deficient cells. Nucleic Acids Res. 2024 Jun 24;52(11):6376–6391. doi: 10.1093/nar/gkae317.PMID: 38721777

Godin SK, Zhang Z, Herken BW, Westmoreland JW, Lee AG, Mihalevic MJ, Yu Z, Sobol RW, Resnick MA, Bernstein KA. The Shu complex promotes error-free tolerance of alkylation-induced base excision repair products. Nucleic Acids Res. 2016 Sep 30;44(17):8199–215. doi: 10.1093/nar/gkw535. Epub 2016 Jun 13.PMID: 27298254

Gonzalez-Huici V, Szakal B, Urulangodi M, Psakhye I, Castellucci F, Menolfi D, Rajakumara E, Fumasoni M, Bermejo R, Jentsch S, Branzei D. DNA bending facilitates the error-free DNA damage tolerance pathway and upholds genome integrity. EMBO J 2014; 33: 327–340 [PMID: 24473148 DOI: 10.1002/embj.201387425]

Gritenaite D, Princz LN, Szakal B, Bantele SC, Wendeler L, Schilbach S, Habermann BH, Matos J, Lisby M, Branzei D, Pfander B. A cell cycle-regulated Slx4-Dpb11 complex promotes the resolution of DNA repair intermediates linked to stalled replication. Genes Dev. 2014 Jul 15;28(14):1604–19. doi: 10.1101/gad.240515.114.PMID: 25030699

Hammond-Martel I, Verreault A, Wurtele H. Chromatin dynamics and DNA replication roadblocks. DNA Repair (Amst). 2021 Aug;104:103140. doi: 10.1016/j.dnarep.2021.103140. Epub 2021 May 23.PMID: 34087728

Hale A, Dhoonmoon A, Straka J, Nicolae CM, Moldovan GL. Multi-step processing of replication stress-derived nascent strand DNA gaps by MRE11 and EXO1 nucleases. Nat Commun. 2023 Oct 7;14(1):6265. doi: 10.1038/s41467-023-42011-0.PMID: 37805499

Han J, Zhang H, Zhang H, Wang Z, Zhou H, Zhang Z. A Cul4 E3 ubiquitin ligase regulates histone hand-off during nucleosome assembly. Cell. 2013 Nov 7;155(4):817–29. doi: 10.1016/j.cell.2013.10.014.PMID: 24209620

Hepp MI, Smolle M, Gidi C, Amigo R, Valenzuela N, Arriagada A, Maureira A, Gogol MM, Torrejón M, Workman JL, Gutiérrez JL. Role of Nhp6 and Hmo1 in SWI/SNF occupancy and nucleosome landscape at gene regulatory regions. Biochim Biophys Acta Gene Regul Mech 2017; 1860: 316–326 [PMID: 28089519 DOI: 10.1016/j.bbagrm.2017.01.002]

Hergeth SP, Schneider R. The H1 linker histones: multifunctional proteins beyond the nucleosomal core particle. EMBO Rep. 2015 Nov;16(11):1439–53. doi: 10.15252/embr.201540749. Epub 2015 Oct 15.

Javaheri A, Wysocki R, Jobin-Robitaille O, Altaf M, Côté J, Kron SJ. Yeast G1 DNA damage checkpoint regulation by H2A phosphorylation is independent of chromatin remodeling. Proc Natl Acad Sci U S A. 2006 Sep 12;103(37):13771–6. doi: 10.1073/pnas.0511192103. Epub 2006 Aug 29.PMID: 16940359

Kalocsay M, Hiller NJ, Jentsch S. Chromosome-wide Rad51 spreading and SUMO-H2A.Z dependent chromosome fixation in response to a persistent DNA double-strand break. Mol Cell. 2009 Feb 13;33(3):335–43. doi:10.1016/j.molcel.2009.01.016.PMID: 19217407

Kamau E, Bauerle KT, Grove A. The Saccharomyces cerevisiae high mobility group box protein HMO1 contains two functional DNA binding domains. J Biol Chem 2004; 279: 55234–55240 [PMID: 15507436 DOI: 10.1074/jbc.M409459200]

Kobor MS, Venkatasubrahmanyam S, Meneghini MD, Gin JW, Jennings JL, Link AJ, Madhani HD, Rine J. A protein complex containing the conserved Swi2/Snf2-related ATPase Swr1p deposits histone variant H2A.Z into euchromatin. PLoS Biol. 2004 May;2(5):E131. doi: 10.1371/journal.pbio.0020131. Epub 2004 Mar 23.PMID: 15045029

Lu J, Kobayashi R, Brill SJ. Characterization of a high mobility group 1/2 homolog in yeast. J Biol Chem 1996; 271: 33678–33685 [PMID: 8969238 DOI: 10.1074/jbc.271.52.33678]

Luger, K., A. W. Mäder, R. K. Richmond, D. F. Sargent and T.J. Richmond, 1997 Crystal structure of the nucleosome core particle at 2.8 A resolution. Nature 389: 251–260.

Malinina DK, Sivkina AL, Korovina AN, McCullough LL, Formosa T, Kirpichnikov MP, Studitsky VM, Feofanov AV. Hmo1 Protein Affects the Nucleosome Structure and Supports the Nucleosome Reorganization Activity of Yeast FACT. Cells 2022; 11 [PMID: 36230893 DOI: 10.3390/cells11192931]

Martino J, Bernstein KA. The Shu complex is a conserved regulator of homologous recombination. FEMS Yeast Res. 2016 Sep;16(6):fow073. doi: 10.1093/femsyr/fow073. Epub 2016 Sep 1.PMID: 27589940

Martire S, Banaszynski LA. The roles of histone variants in fine-tuning chromatin organization and function. Nat Rev Mol Cell Biol. 2020 Sep;21(9):522–541. doi: 10.1038/s41580-020-0262-8. Epub 2020 Jul 14.PMID: 32665685

McCauley MJ, Huo R, Becker N, Holte MN, Muthurajan UM, Rouzina I, Luger K, Maher LJ 3rd, Israeloff NE, Williams MC. Single and double box HMGB proteins differentially destabilize nucleosomes. Nucleic Acids Res 2019; 47: 666–678 [PMID: 30445475 DOI: 10.1093/nar/gky1119]

Mejia-Ramirez E, Limbo O, Langerak P, Russell P. Critical function of gH2A in S-phase. PLoS Genet 2015; 11:e1005517; 10.1371/journal.pgen.1005517

Meng X, Zhao X. Replication fork regression and its regulation. FEMS Yeast Res. 2017 Jan 1;17(1):fow110. doi: 10.1093/femsyr/fow110.PMID: 28011905

Mizuguchi G, Shen X, Landry J, Wu WH, Sen S, Wu C. ATP-driven exchange of histone H2AZ variant catalyzed by SWR1 chromatin remodeling complex. Science. 2004 Jan 16;303(5656):343–8. doi: 10.1126/science.1090701. Epub 2003 Nov 26.PMID: 14645854

Moldovan GL, Pfander B, Jentsch S. PCNA, the maestro of the replication fork Cell. 2007 May 18;129(4):665–79. doi: 10.1016/j.cell.2007.05.003.PMID: 17512402

Ohouo PY, Bastos de Oliveira FM, Almeida BS, Smolka MB. 2010. DNA damage signaling recruits the Rtt107–Slx4 scaffolds via Dpb11 to mediate replication stress response. Mol Cell 39: 300–306.

Ohouo PY, Bastos de Oliveira FM, Liu Y, Ma CJ, Smolka MB. DNA-repair scaffolds dampen checkpoint signalling by counteracting the adaptor Rad9. Nature. 2013; 493:120–4. doi: 10.1038/nature11658.

Panday A, Grove A. The high mobility group protein HMO1 functions as a linker histone in yeast. Epigenetics Chromatin 2016; 9: 13 [PMID: 27030801 DOI: 10.1186/s13072-016-0062-8]

Panday A, Grove A. Yeast HMO1: Linker Histone Reinvented. Microbiol Mol Biol Rev 2017; 81 [PMID: 27903656 DOI: 10.1128/MMBR.00037-16]

Panday A, Xiao L, Grove A. Yeast high mobility group protein HMO1 stabilizes chromatin and is evicted during repair of DNA double strand breaks. Nucleic Acids Res 2015; 43: 5759–5770 [PMID: 25979266 DOI: 10.1093/nar/gkv498]

Panday A, Xiao L, Gupta A, Grove A. Control of DNA end resection by yeast Hmo1p affects efficiency of DNA end-joining. DNA Repair (Amst). 2017 May;53:15–23. doi: 10.1016/j.dnarep.2017.03.002. Epub 2017 Mar 9.PMID: 28336179

Papamichos-Chronakis M, Watanabe S, Rando OJ, Peterson CL. Global regulation of H2A.Z localization by the INO80 chromatin-remodeling enzyme is essential for genome integrity. Cell. 2011 Jan 21;144(2):200–13. doi: 10.1016/j.cell.2010.12.021.PMID: 21241891

Pardo B, Crabbé L, Pasero P. Signaling pathways of replication stress in yeast. FEMS Yeast Res. 2017 Mar 1;17(2). doi: 10.1093/femsyr/fow101.PMID: 27915243

Pham P, Wood EA, Cox MM, Goodman MF. RecA and SSB genome-wide distribution in ssDNA gaps and ends in Escherichia coli. Nucleic Acids Res. 2023 Jun 23;51(11):5527–5546. doi: 10.1093/nar/gkad263.PMID: 37070184

Postnikov YV, Bustin M. Functional interplay between histone H1 and HMG proteins in chromatin. Biochim Biophys Acta. 2016 Mar;1859(3):462–7. doi: 10.1016/j.bbagrm.2015.10.006. Epub 2015 Oct 8.

Serra-Cardona A, Hua X, McNutt SW, Zhou H, Toda T, Jia S, Chu F, Zhang Z. The PCNA-Pol δ complex couples lagging strand DNA synthesis to parental histone transfer for epigenetic inheritance. Sci Adv. 2024 Jun 7;10(23):eadn5175. doi: 10.1126/sciadv.adn5175. Epub 2024 Jun 5.PMID: 38838138

Serra-Cardona A, Zhang Z. Replication-Coupled Nucleosome Assembly in the Passage of Epigenetic Information and Cell Identity. Trends Biochem Sci. 2018 Feb;43(2):136–148. doi: 10.1016/j.tibs.2017.12.003. Epub 2017 Dec 29.PMID: 29292063

Sherman F. Getting started with yeast. Methods Enzymol. 2002;350:3–41. doi: 10.1016/s0076-6879(02)50954-x.PMID: 12073320

Shi G, Yang C, Wu J, Lei Y, Hu J, Feng J, Li Q. DNA polymerase δ subunit Pol32 binds histone H3-H4 and couples nucleosome assembly with Okazaki fragment processing. Sci Adv. 2024 Aug 9;10(32):eado1739. doi: 10.1126/sciadv.ado1739. Epub 2024 Aug 9.PMID: 3912122

Siler J, Guo N, Liu Z, Qin Y, Bi X. γH2A/γH2AX Mediates DNA Damage-Specific Control of Checkpoint Signaling in *Saccharomyces cerevisiae*. Int J Mol Sci. 2024 Feb 20;25(5):2462. doi: 10.3390/ijms25052462.PMID: 38473708

Simpson, R.T., F. Thoma, and J. M. Brubaker, 1985 Chromatin reconstituted from tandemly repeated cloned DNA fragments and core histones: a model system for study of higher order structure. Cell 42: 799–808.

Sogo JM, Lopes M, Foiani M. Fork reversal and ssDNA accumulation at stalled replication forks owing to checkpoint defects. Science. 2002; 297: 599–602. PMID: 12142537

Stros M. HMGB proteins: interactions with DNA and chromatin. Biochim Biophys Acta 2010; 1799: 101–113 [PMID: 20123072 DOI: 10.1016/j.bbagrm.2009.09.008]

Srivatsan A, Li BZ, Szakal B, Branzei D, Putnam CD, Kolodner RD. The Swr1 chromatin remodeling complex prevents genome instability induced by replication fork progression defects. Nat Commun. 2018 Sep 11;9(1):3680. doi: 10.1038/s41467-018-06131-2.PMID: 30206225

Szakal B, Branzei D. Premature Cdk1/Cdc5/Mus81 pathway activation induces aberrant replication and deleterious crossover. EMBO J. 2013 Apr 17;32(8):1155–67. doi: 10.1038/emboj.2013.67. Epub 2013 Mar 26.PMID: 23531881

Tian C, Zhang Q, Jia J, Zhou J, Zhang Z, Karri S, Jiang J, Dickinson Q, Yao Y, Tang X, Huang Y, Guo T, He Z, Liu Z, Gao Y, Yang X, Wu Y, Chan KM, Zhang D, Han J, Yu C, Gan H. DNA polymerase delta governs parental histone transfer to DNA replication lagging strand. Proc Natl Acad Sci U S A. 2024 May 14;121(20):e2400610121. doi: 10.1073/pnas.2400610121. Epub 2024 May 7.PMID: 3871362

Toda T, Fang Y, Shan CM, Hua X, Kim JK, Tang LC, Jovanovic M, Tong L, Qiao F, Zhang Z, Jia S. Mrc1 regulates parental histone segregation and heterochromatin inheritance. Mol Cell. 2024 Sep 5;84(17):3223–3236.e4. doi: 10.1016/j.molcel.2024.07.002. Epub 2024 Aug 1.PMID: 39094566

Tsabar M, Eapen VV, Mason JM, Memisoglu G, Waterman DP, Long MJ, Bishop DK, Haber JE. Caffeine impairs resection during DNA break repair by reducing the levels of nucleases Sae2 and Dna2. Nucleic Acids Res. 2015 Aug 18;43(14):6889–901. doi: 10.1093/nar/gkv520. Epub 2015 May 27.PMID: 26019182

Williams JS, Williams RS, Dovey CL, Guenther G, Tainer JA, Russell P. γH2A binds Brc1 to maintain genome integrity during S-phase EMBO J. 2010 Mar 17;29(6):1136–48. doi: 10.1038/emboj.2009.413. Epub 2010 Jan 21.PMID: 20094029

Xiao L, Williams AM, Grove A. The C-terminal domain of yeast high mobility group protein HMO1 mediates lateral protein accretion and in-phase DNA bending. Biochemistry 2010; 49: 4051–4059 [PMID: 20402481 DOI: 10.1021/bi1003603]

Yu C, Gan H, Serra-Cardona A, Zhang L, Gan S, Sharma S, Johansson E, Chabes A, Xu RM, Zhang Z. A mechanism for preventing asymmetric histone segregation onto replicating DNA strands. Science. 2018 Sep 28;361(6409):1386–1389. doi: 10.1126/science.aat8849. Epub 2018 Aug 16.PMID: 30115745

Yu J, Zhang Y, Fang Y, Paulo JA, Yaghoubi D, Hua X, Shipkovenska G, Toda T, Zhang Z, Gygi SP, Jia S, Li Q, Moazed D. A replisome-associated histone H3-H4 chaperone required for epigenetic inheritance. Cell. 2024 Sep 5;187(18):5010–5028.e24. doi: 10.1016/j.cell.2024.07.006. Epub 2024 Aug 1.PMID: 39094570

Yu Y, Srinivasan M, Nakanishi S, Leatherwood J, Shilatifard A, Sternglanz R. A conserved patch near the C terminus of histone H4 is required for genome stability in budding yeast Mol Cell Biol. 2011 Jun;31(11):2311–25. doi: 10.1128/MCB.01432-10. Epub 2011 Mar 28.PMID: 21444721

Zhang W, Feng J, Li Q. The replisome guides nucleosome assembly during DNA replication. Cell Biosci. 2020 Mar 12;10:37. doi: 10.1186/s13578-020-00398-z. eCollection 2020.PMID: 32190287

Zhang Z, Shibahara K, Stillman B. PCNA connects DNA replication to epigenetic inheritance in yeast. Nature. 2000 Nov 9;408(6809):221–5. doi: 10.1038/35041601.PMID: 11089978

